# Temporal Phosphoproteomic Network Reconstruction of Condition-Dependent EGFR Signaling Routes Through Unmeasured Proteins

**DOI:** 10.64898/2026.03.20.712775

**Authors:** Heebok Lee, Gihoon Lee

## Abstract

Time-resolved phosphoproteomics captures fast signaling dynamics, but proteins that show no phosphorylation change, or escape detection, leave gaps the data cannot bridge. TS-BFS-BS uses the temporal order of a phosphoproteomic time course to guide a beam-pruned breadth-first search over STRING v12.5 functional, physical, and regulatory priors, returning temporally continuous routes that can pass through unmeasured bridge proteins. In EGF time courses from MDA-MB-468, EGFR Flp-In HEK293T and HeLa cells, endpoints and short routes were stable across search settings, and endpoints were largely shared across STRING priors, whereas long routes depended on settings and prior. When measured hubs CBL and SRC were removed, both were recovered as bridges and 94–99% of endpoints were retained. Incorrect temporal orders gave endpoint Jaccard indices of only 0.23–0.33 with the correct order, comparable to SHP2 inhibition (0.25–0.35), under which PTPN11 was excluded from all routes by its measured pretreatment response and returned after washout. TS-BFS-BS generates time-resolved, testable signaling hypotheses for iterative refinement.

## Introduction

Mass spectrometry-based phosphoproteomics quantifies thousands of phosphorylation sites, and time-resolved sampling can follow how a signal spreads through the cell after a stimulus^1–3^, including the oncogenic signals that drive disease progression^4^. Its view of the signaling network is nevertheless incomplete^5^. Many proteins that relay signals between the detected ones remain invisible because they carry no regulated site, change too little to pass a response threshold, or yield peptides that are not detected. Moreover, consecutive changes in phosphorylation do not prove a direct interaction, and several reactions may occur within a single sampling interval^6^. Signaling routes inferred from phosphoproteomic data alone are therefore fragmented, and prior knowledge of protein interactions is needed to connect them. STRING, for example, assigns each known association a confidence score that combines evidence from experiments, curated pathway databases, co-expression, and text mining^7^.

Such prior knowledge can bridge missing measurements, but it introduces uncertainties of its own. Static databases list interactions that are possible in general, not those that are active in a given cell and time window^8^. Highly connected proteins cause the number of candidate paths to grow combinatorially, whereas aggressive pruning may remove intermediates that a later target requires^9^. Likewise, restricting the search to measured proteins creates artificial gaps, while admitting unmeasured proteins without limit produces nonspecific routes. A useful method must therefore constrain the prior network without losing temporally continuous and biologically plausible alternatives. Existing approaches, including constraint-based pathway synthesis^6^, network propagation^9^, and graph-based prior knowledge networks and causal reasoning^10–12^, each address parts of this problem. Where timing is used, it typically orders a single event per protein, such as its first significant change^6^, rather than letting every sampling interval constrain which routes can continue into the next.

We developed TS-BFS-BS (Temporal-Sequential Breadth-First Search and Beam Search), an algorithm that reconstructs candidate signaling routes from time-resolved phosphoproteomic data by searching a STRING interaction network^7^. Its name reflects its four components. The time course is divided into consecutive response intervals, and each protein’s response is assessed separately within each interval (temporal). The search proceeds interval by interval: the measured proteins reached in one interval become the starting points of the next, so that every route follows the order of the measurements (sequential). Within each interval, a breadth-first search connects these starting points to the proteins that respond in that interval, allowing at most a small, fixed number of intermediates that do not respond in that interval, whether unmeasured or measured without a response (breadth-first search). Throughout, the search uses only high-confidence interactions and retains the best-supported candidates at each step, while keeping at least one route for every protein reached (beam search). The functional, physical, and regulatory networks of STRING are treated as distinct hypothesis spaces rather than interchangeable versions of one network^7^. The result is an ensemble of temporally consistent, STRING-supported candidate routes that can pass through proteins the experiment did not observe. Intermediate proteins are excluded on cell-context grounds only when expression data for the cell line are explicitly negative^13^.

TS-BFS-BS was designed to keep manual intervention to a minimum. Measured phosphorylation data are processed automatically by general rules that apply equally to every protein; no protein is added, removed, or reweighted by hand, and the same settings are used for every dataset. We kept the algorithm deliberately simple so that its behavior can be traced to the data and the STRING prior, and so that it can later be combined with other models and experimental data types.

We applied TS-BFS-BS to signaling by the EGF receptor (EGFR), a well-characterized system that is temporally organized and context-dependent rather than a fixed cascade^14^. After receptor stimulation, phosphorylation, adaptor recruitment, cytoskeletal remodeling, metabolic regulation, transcriptional control, and feedback unfold over distinct but overlapping timescales^14^, and published time courses are available for several conditions and cell lines^1,6,15^. Using an EGF time course of MDA-MB-468 breast cancer cells that includes untreated, SHP2-inhibited, and inhibitor-washout conditions^15^, we compared reconstructions under the three STRING priors, tested whether the method recovers measured proteins deleted from the input, asked whether the temporal order of the measurements shapes the result, and reconstructed the inhibitor and washout conditions. We then applied the same settings to EGF time courses from EGFR Flp-In HEK293T^6^ and HeLa cells^1^ and compared TS-BFS-BS with the Temporal Pathway Synthesizer (TPS) on the dataset used in its original description^6^. Together, these analyses show what the method reconstructs, which parts of its output are robust, and how its results depend on the STRING prior, the temporal order, and the cellular context.

## Results

### TS-BFS-BS reconstructs temporally continuous routes from interval-wise responses

TS-BFS-BS takes three inputs: a phosphoproteomic time course, a stimulus source (here EGFR), and a STRING v12.5 interaction network^7^. From these, it returns an ensemble of candidate signaling routes (Fig. 1).

**Figure 1.**
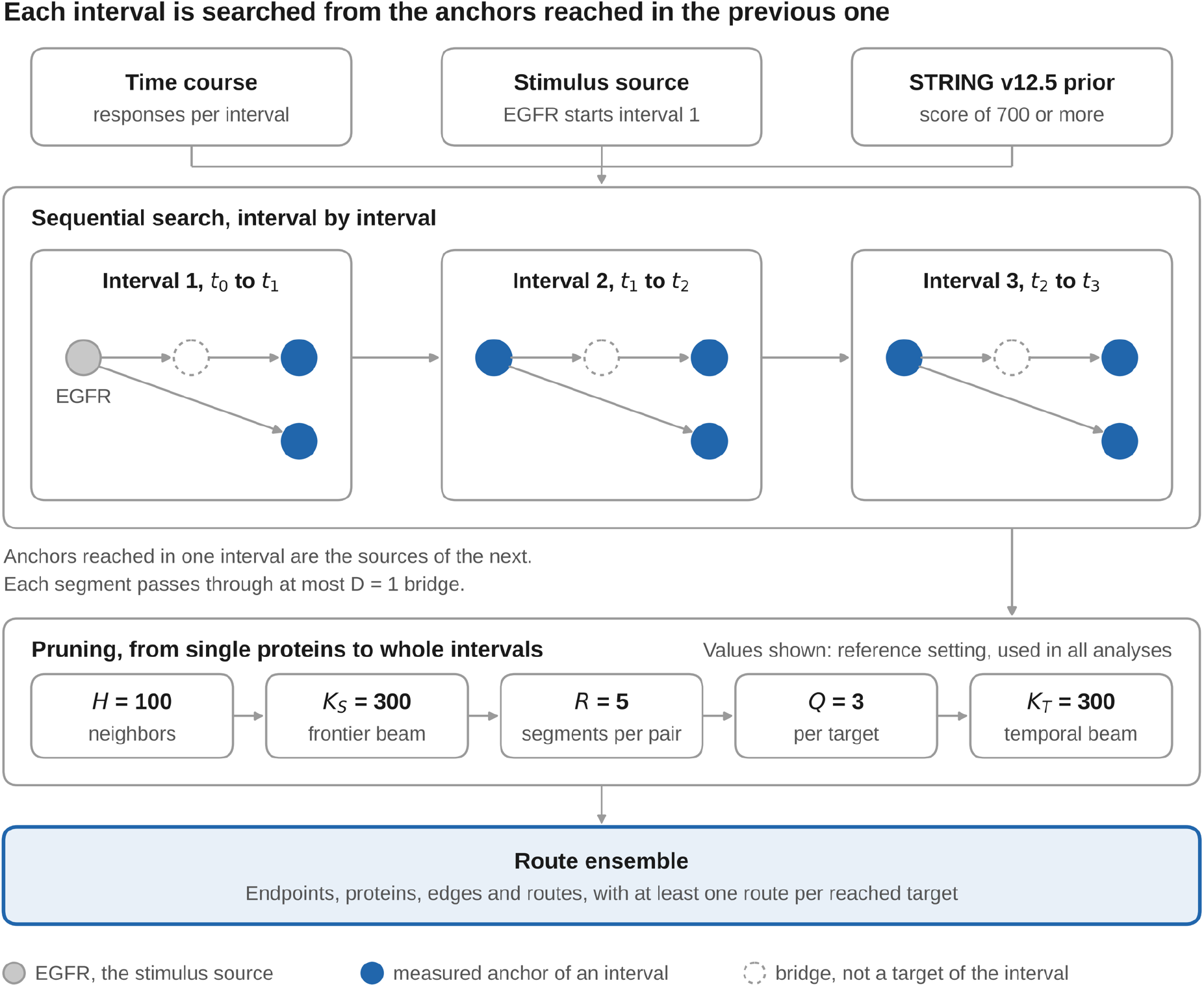
Overview of TS-BFS-BS. The three inputs, a phosphoproteomic time course, the stimulus source EGFR, and a STRING v12.5 interaction network, feed a search that proceeds interval by interval. In each interval, segments, each consisting of one or two edges of the STRING prior network, link a source to the measured anchors of that interval, either directly or through at most one bridge, a protein that is not an anchor of the interval. The anchors reached become the sources of the next interval. Five nested controls prune the search, and at least one route is kept for every reached anchor. The schematic shows the logic of the search; the nodes are not specific proteins. The values shown are the settings used in all analyses after the parameter study (Table 1).

**Table 1.**
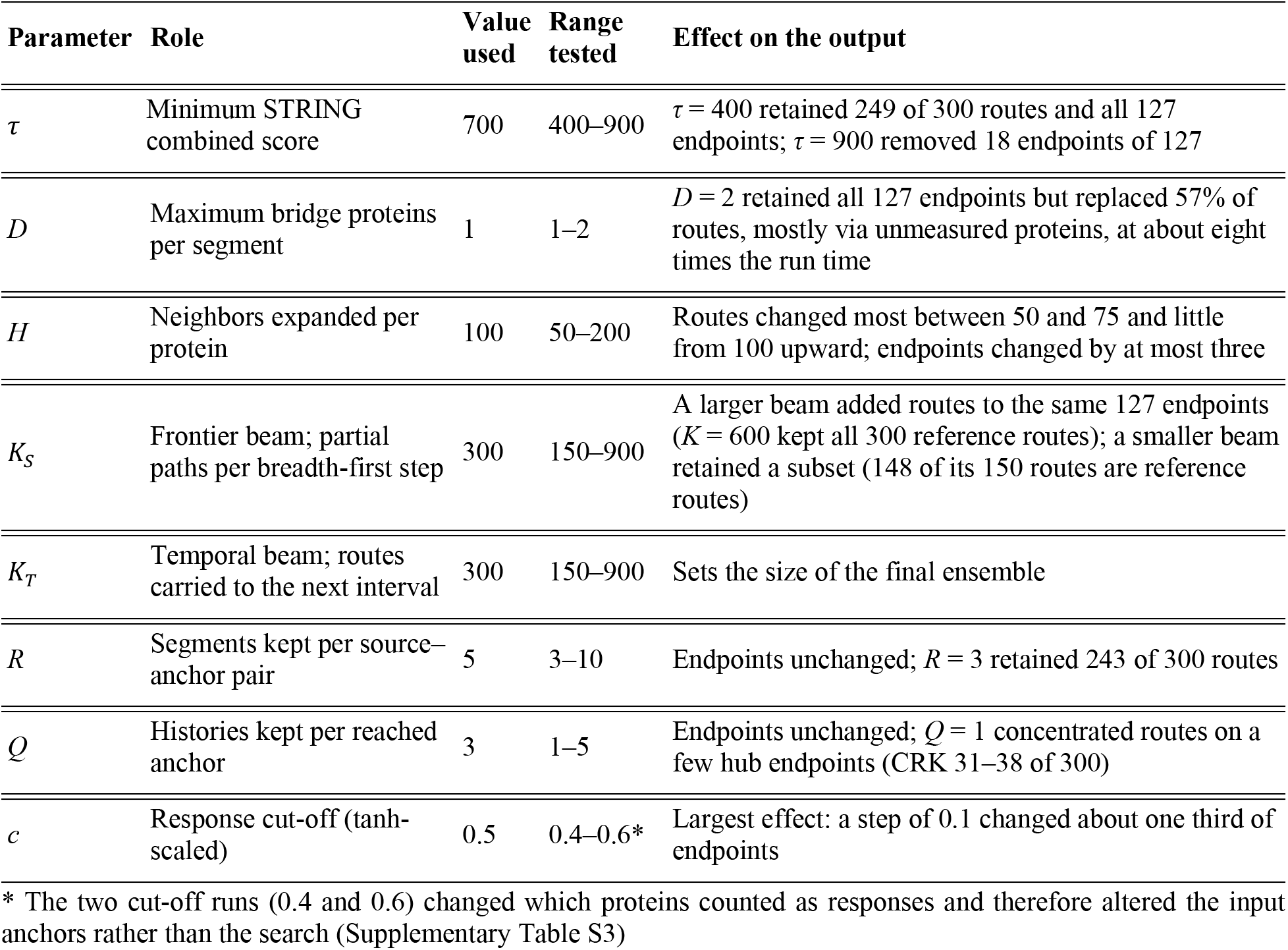
Parameters of TS-BFS-BS, the values used in this study and their effect on the output. Effects were measured on the EGF-stimulated MDA-MB-468 time course under the functional STRING prior. All runs were compared with a reference run at *H* = 100. Full results are given in Supplementary Table S3 and Supplementary Fig. S1.

The time course is first divided into consecutive intervals. Within each interval, a protein is called a response anchor if its phosphorylation changed beyond a fixed cut-off *c* (tanh-scaled; Table 1) during that interval. Anchors are selected by the magnitude of the change alone. Whether a site is activating or inhibitory is taken from a curated table of regulatory sites and, for the remaining sites, from OmniPath^16^ and used only for eligibility: a gene with an opposing and no supporting site in an interval is not used as a target or bridge in that interval (Methods).

The search then proceeds interval by interval. It starts from EGFR in the first interval and, in each later interval, from the anchors reached in the previous one, so that every route passes through measured responses in the order in which they were observed. Here, “route” always means a complete route: one that starts at EGFR, passes through every interval and ends on an anchor of the last interval. The endpoint of a route is its last protein. Because several routes can end on the same protein, the number of endpoints in an ensemble, counted as distinct proteins, is typically smaller than the number of routes. Within an interval, a breadth-first search links each starting protein to each anchor of that interval; each such link is a segment. A segment may pass through at most *D* intermediate proteins that are not anchors of the interval, whether they were never measured or were measured without a response. These bridges allow a route to continue across proteins that the experiment did not observe, and their number is bounded so that the search cannot connect any two proteins through arbitrary hubs.

Five nested controls keep the search tractable while retaining alternative routes. A degree cap *H* limits the neighbors expanded per protein, and a frontier beam *K_S_* limits the partial paths kept at each breadth-first step within an interval. A cap *R* limits the segments kept per source–anchor pair, and a quota *Q* limits the number of distinct upstream route prefixes (histories) kept per reached anchor. Finally, a temporal beam *K_T_* limits the total routes carried forward to the next interval. Because at least one route is kept for every anchor reached, pruning removes alternative routes but never a reached protein. Candidates are ranked by the product of the STRING combined scores of their edges, so that a route ranks higher when it has fewer or higher-scoring edges; these scores order the candidates but are not interpreted as probabilities.

We next asked which parts of the output depend on these settings. Starting from a reference setting (*τ* = 700*, D* = 1, *K_S_* = *K_T_* = 300, *c* = 0.5), we varied one parameter at a time, and *R* and *Q* jointly, on the EGF-stimulated MDA-MB-468 time course^15^ under the functional STRING prior. Each run was compared with its reference run by the Jaccard similarity of four sets: complete routes, proteins, directed edges, and endpoints (Fig. 2, Supplementary Fig. S1 and Supplementary Table S3). All runs were compared with a reference run at *H* = 100, the setting adopted for all subsequent analyses. The search is deterministic: an independent repeat of the reference setting returned an identical set of 300 routes. The parameters and their values are listed in Table 1. The full algorithm is described in Methods.

**Figure 2.**
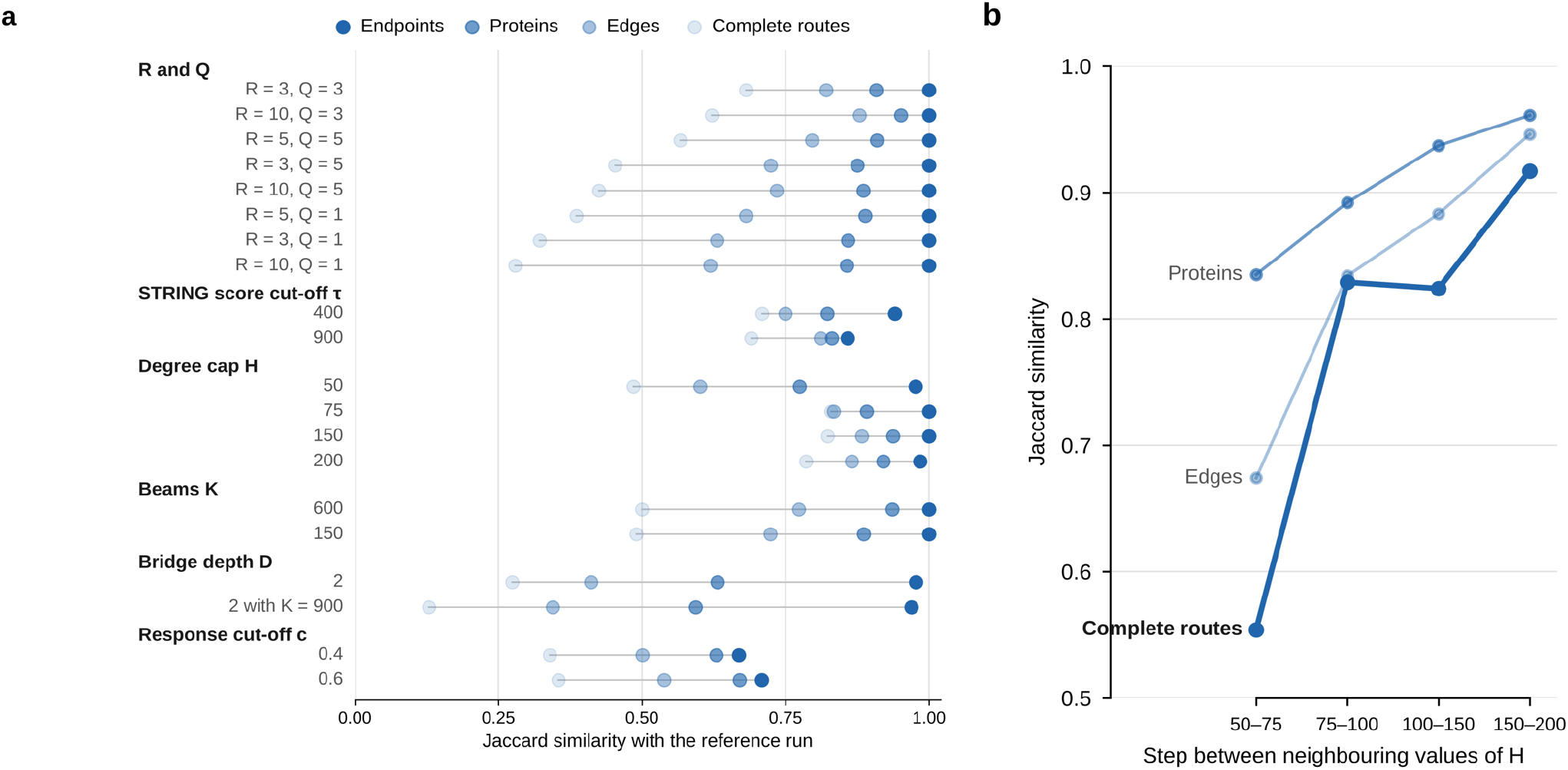
Sensitivity of TS-BFS-BS output to search parameters. **(a)** Similarity of each parameter run to the reference run. Each row is one run, grouped by the parameter changed; the four dots give the Jaccard similarity of its endpoints, proteins, directed edges, and complete routes to those of the reference run (*τ* = 700, *D* = 1, *K_S_ = K_T_ =* 300, *R* = 5, *Q* = 3, *c* = 0.5; *H* = 100). Endpoints changed least and complete routes most. **(b)** Effect of the degree cap *H*. Jaccard similarity of proteins, directed edges and complete routes between runs at adjacent values of *H* (50→75, 75→100, 100→150 and 150→200). Of the 300 routes at the lower value, 214, 272, 271, and 287, respectively, were retained at the higher value. Route content changed most between *H* = 50 and 75 and little from *H* = 100 upward.

### The three STRING priors agree on endpoints but explain them differently

We reconstructed signaling routes in EGF-stimulated MDA-MB-468 cells under the functional, physical, and regulatory STRING priors, using the same input, settings, and cell-context filter^13^. Each run returned 300 routes, the limit set by the temporal beam *K_T_*, so route number carries no biological information. The comparison therefore rests on the content of each ensemble (Table 2). For every pair of STRING priors, agreement was highest for endpoints and decreased through proteins and edges to routes. The functional and physical STRING priors reached nearly the same endpoints (118 of 127 shared; Jaccard 0.93) but shared only 132 of 468 distinct routes (Jaccard 0.28). For 51 of the 118 shared endpoints, the two STRING priors had no route in common, so the same response was reached by a different path. All 34 regulatory endpoints were also reached under both undirected STRING priors, yet the regulatory routes shared almost nothing with them (8 identical routes with each; Jaccard 0.01) (Supplementary Fig. S2a and Supplementary Table S5). The endpoints are thus set largely by the data, and the routes by the STRING prior.

**Table 2.** Route ensembles of the EGF-stimulated MDA-MB-468 under three STRING priors.

|  | Functional | Physical | Regulatory |
|---|---|---|---|
| Proteins, excluding EGFR | 278 | 258 | 94 |
| Directed edges | 396 | 364 | 196 |
| Endpoints | 127 | 118 | 34 |
| Distinct first hops | 39 | 38 | 23 |
| Routes through the three most frequent proteins | 44% (CBL, HSP90AA1, HGS) | 43% (CBL, HSP90AA1, UBC) | 83% (SRC, PTPN11, STAT3) |
| Jaccard with functional: endpoints / exact routes | – | 0.93 / 0.28 | 0.27 / 0.01 |

A small core was nevertheless common to all three STRING priors: 6 identical routes, 21 directed edges and the 34 endpoints (Fig. 3a and Supplementary Table S7). This core is centered on direct EGFR partners, namely CBL, CRK, EPS15, GAB1, GRB2, PIK3R2, PLCG2, SRC and STAT3. Its identical routes include EGFR→SRC→CDCP1, which links the receptor to a Src-family substrate known to cooperate with EGFR in breast cancer cells^17^. These shared elements are the parts of the reconstruction that do not depend on the choice of STRING prior.

**Figure 3.**
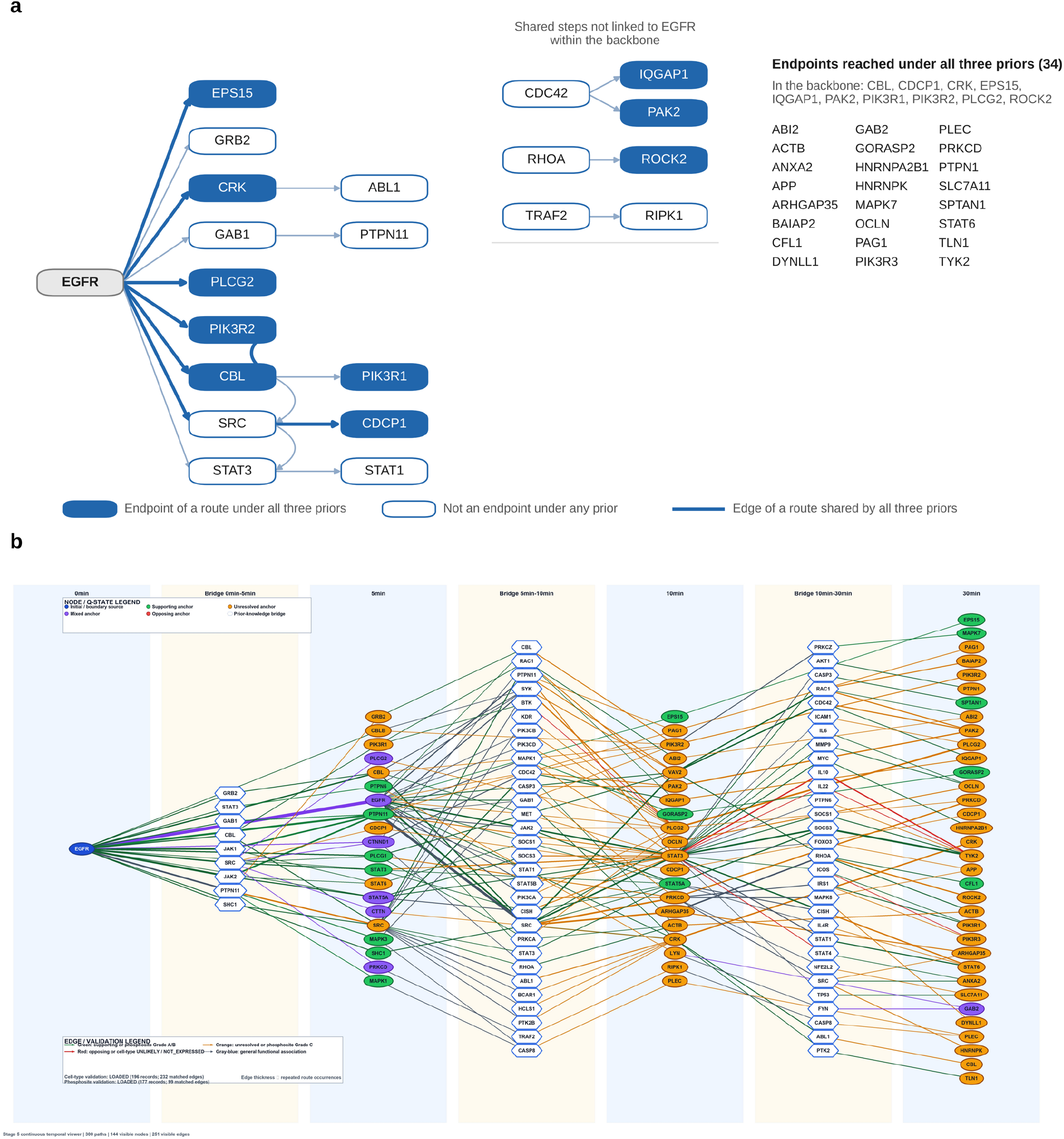
Route reconstructions of EGF-stimulated MDA-MB-468 cells under the three STRING priors. **(a)** Backbone shared by the functional, physical, and regulatory STRING priors: the 21 directed edges present in all three ensembles of 300 routes. Edges are pooled across routes, so the backbone is not itself a route. Filled nodes are proteins at which at least one route ends in each of the three ensembles (34 such endpoints in total; those not in the backbone are listed on the right); open nodes are not endpoints in all three ensembles. Because a protein can be the endpoint of some routes and an intermediate in others, filled nodes may have outgoing edges. Thick edges belong to the six complete routes common to all three STRING priors. **(b)** Interval-resolved map of the regulatory ensemble (300 routes; 144 nodes and 251 edges). Columns alternate between measured time points and the bridge layer of each interval. Ellipses are measured anchors, colored by the polarity of the segments that end on them in that interval: supporting (green) or opposing (red) the observed direction of change, unresolved (orange) or mixed (purple) when these segments carry more than one polarity. Hexagons are bridges supplied by the STRING prior. Edge colors follow the rules in Methods, and edge width scales with the number of routes using the edge. The route network and route maps for each STRING prior are shown in Supplementary Figs. S3–S5.

The functional STRING prior gave the broadest view. Its ensemble was organized around ubiquitin-dependent sorting of the activated receptor: about two-fifths of its routes left EGFR through ubiquitin and endocytic proteins, tracing CBL→ubiquitin (UBC)→EPS15, HGS and STAM2 and continuing to RAB5–RAB7–RAB11 trafficking^18,19^. It also reconstructed the PAG1–CSK feedback that restrains Src-family kinases^20^. Its additional coverage, however, was its least secure part. Nine endpoints were reached under this STRING prior alone, each by a single route of four to six edges, and about a tenth of the routes ran through abundant chaperones and RNA-binding proteins, whose functional associations need not reflect signaling. The route network for the functional STRING prior is shown in Supplementary Fig. S3.

The physical STRING prior gave shorter and more specific routes. Its ensemble reached a subset of the functional endpoints, with no endpoint of its own, and often did so through shorter routes that stayed within known complexes; fewer of its routes reached six edges (59, against 81 under the functional STRING prior; Supplementary Fig. S2b). For example, EGFR→RASA1→ARHGAP35 replaced a four-edge functional route and corresponds to the p120RasGAP–p190RhoGAP complex formed after growth factor stimulation^21^, and EGFR→CTNNA1→TJP1→OCLN remained within junctional proteins. Because physical edges also record ubiquitin conjugation, ubiquitin acted as a hub in 12% of routes; such edges should not be read as steps in a signal relay. The route network for the physical STRING prior is shown in Supplementary Fig. S4.

The regulatory STRING prior gave a narrow, directed view. Its ensemble was small and concentrated: most routes passed through SRC, PTPN11, or STAT3, and JAK–STAT proteins appeared in half of them, compared with 4–5% under the undirected STRING priors. The route network and the route map for the regulatory STRING prior are shown in Fig. 3b and Supplementary Fig. S5.

Directionality allowed several routes to be read as regulatory chains, such as EGFR→SRC→ARHGAP35→RHOA→ROCK2^22,23^. PTPN11 (SHP2) was prominent only under this STRING prior (118 routes, against 4 and 1 under the functional and physical STRING priors), which makes it the most informative STRING prior for the SHP2-inhibitor analysis below. Two features limit its interpretation. First, 29% of its routes crossed intermediates whose action requires new gene expression or secretion, such as SOCS3, CISH and IL10, which are unlikely to act within 30 min of stimulation^24^. Second, as with the total route number, the number of routes per endpoint reflects how the beam was filled; the 54 routes ending at TYK2, for example, are alternative paths to one endpoint, not stronger evidence (Supplementary Fig. S2c).

Under all three STRING priors, the cell-context filter admitted some bridges that are unexpected in this cell line. These include PTEN, which carries a homozygous inactivating mutation in MDA-MB-468 cells^25^, and ESR1, which is not expected in a triple-negative line; each appeared on only a few routes. Because the filter excludes a protein only when both RNA and protein data are negative, such bridges are retained as unverified hypotheses.

Together, these results show that the choice of STRING prior determines the explanation more than the set of responses explained. We therefore report results that hold under all three STRING priors as the most robust and attribute the rest to the semantics of the STRING prior that produced them.

### Proteins removed from the input are recovered as bridges under all three STRING priors

TS-BFS-BS was designed to reconstruct routes through proteins that the experiment did not observe. To test this directly, we deleted two measured proteins from the EGF-stimulated MDA-MB-468 input and asked whether the search would recover them.

We chose CBL and SRC because they are the only measured proteins that occur as intermediates in at least 30 of the 300 reference routes under each of the three STRING priors, so deleting them tests recovery under every STRING prior rather than under one. All three of their phosphosite entries (CBL Y141 and SRC Y193 and Y222) were removed from the pre-stimulation reference and from every interval, and nothing else was changed (958 to 955 entries). The modified input thus describes an experiment in which neither protein was detected. Once unmeasured, a protein can no longer be an anchor or an endpoint, but it can still enter a route as a bridge between two measured anchors.

The reconstruction was largely preserved (Fig. 4 and Table 3). Endpoint sets were retained almost completely: 126 of 127, 117 of 118 and 32 of 34 under the functional, physical and regulatory STRING priors (Jaccard 0.99, 0.99 and 0.94). The endpoints lost were CBL itself, which cannot be an endpoint once unmeasured, and, under the regulatory STRING prior, CDCP1. Routes that had used neither protein were nearly all reproduced (200 of 216, 210 of 214 and 134 of 134).

**Figure 4.**
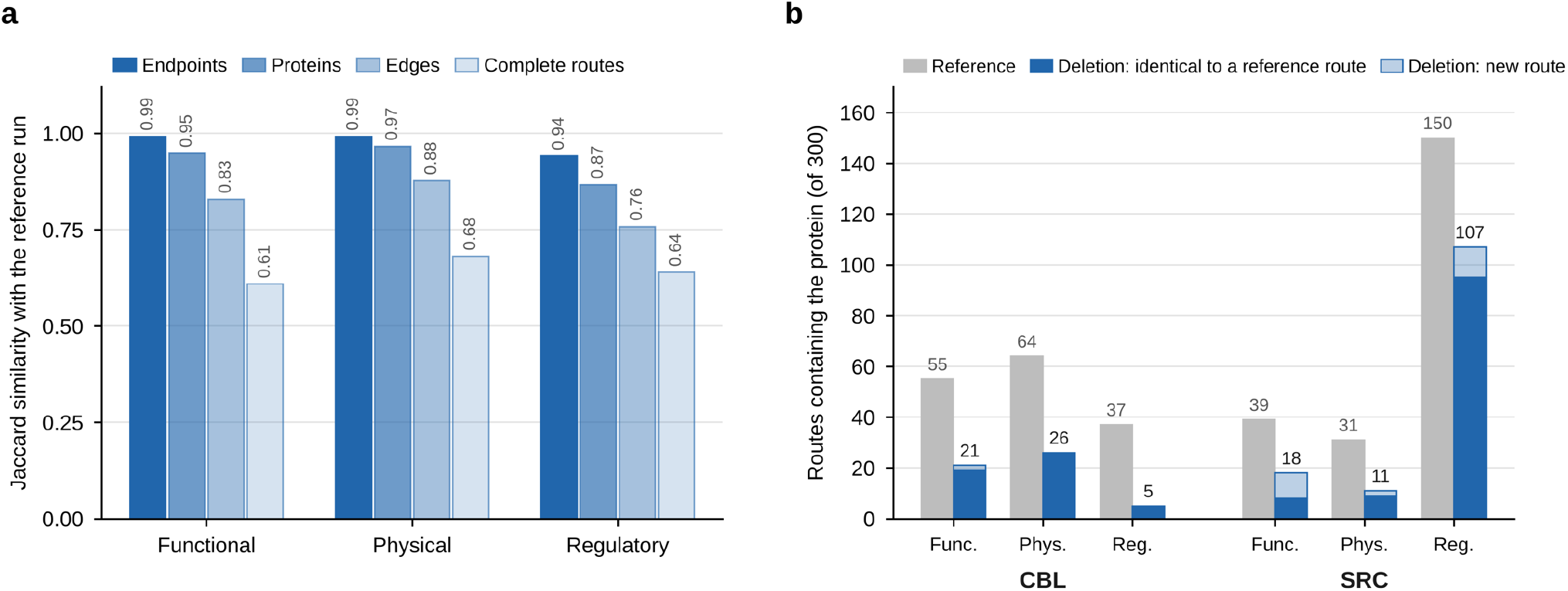
CBL and SRC are recovered as bridges after deletion from the input. CBL and SRC were removed from the EGF-stimulated MDA-MB-468 input, and the search was repeated under each STRING prior with the reference setting. **(a)** Jaccard similarity of endpoints, proteins, directed edges, and complete routes between the deletion run and the reference run under each STRING prior. **(b)** Number of routes (of 300) containing CBL or SRC in the reference run (gray) and in the deletion run (blue). In the deletion run, the two proteins are unmeasured and can enter routes only as bridges. Each deletion bar is divided into routes identical to a reference route (dark blue) and new routes (light blue). See also Table 3 and Supplementary Fig. S7.

**Table 3.**
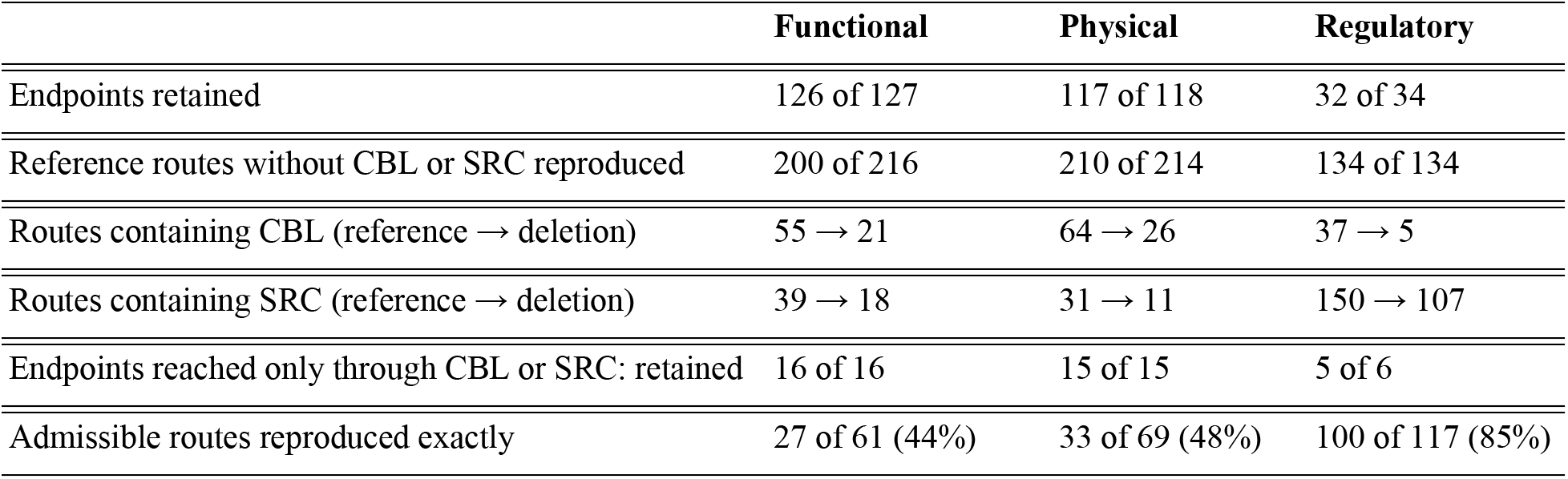
Effect of deleting CBL and SRC from the EGF-stimulated MDA-MB-468 input.

Both proteins returned under every STRING prior: CBL in 21, 26 and 5 routes and SRC in 18, 11 and 107 (functional, physical and regulatory, respectively). As the search rules require, each recovered protein appeared only as a bridge between two measured anchors, and in most routes it returned with the same neighbors it had when observed (Supplementary Table S9). CBL reappeared almost exclusively directly after EGFR (20 of 21 functional and all 26 physical routes), most often followed by UBC (Supplementary Fig. S6). Under the regulatory STRING prior, SRC reappeared mainly after PTPN11 (69 of 107 routes) or after EGFR.

Endpoints that had depended on the deleted proteins were also retained. Under the functional STRING prior, 16 endpoints had been reached only through CBL or SRC; all 16 remained endpoints, 9 of them again reached through the deleted proteins and the other 7 through alternative routes. Those include EGFR→LYN→SRC→CSK, EGFR→STAT3→SRC→PTPN1, EGFR→LYN→SRC→CDCP1 and EGFR→CBL→UBC→ITCH→SCAMP3. The physical and regulatory STRING priors behaved in the same way, retaining all 15 and 5 of 6 such endpoints, respectively; the one lost under the regulatory STRING prior was CDCP1. These placements agree with known biology, although this agreement is not independent evidence, because the STRING prior draws on the same literature: CBL is the E3 ligase that ubiquitinates the activated receptor^18^, CSK is the principal negative regulator of Src-family kinases^26^, CDCP1 is a Src-family substrate^17,27^, and PTP1B (PTPN1) dephosphorylates and activates c-Src in breast cancer cells^28^ (Supplementary Fig. S7 and Supplementary Table S8).

Recovery at the level of whole routes was partial, for two reasons. First, a deleted protein cannot act as an anchor, so routes in which CBL or SRC relayed the signal from one interval to the next could not return (20, 14 and 46 routes under the functional, physical, and regulatory STRING priors). About half of these routes contained CBL directly followed by SRC; with both proteins unmeasured, such a route would require two consecutive bridges, which D = 1 does not allow. Second, as a bridge, a recovered protein must compete with other candidates on STRING prior score. To measure this effect, we restricted the comparison to admissible routes: reference routes containing CBL or SRC that could still be formed under the search rules once both proteins were unmeasured. Such a route must pass through one measured anchor in each interval, may contain CBL or SRC only as a bridge between two anchors, with at most one bridge per segment, and must not end on either protein. For example, EGFR→CBL→GRB2 is admissible, because CBL can bridge EGFR and the measured GRB2, whereas a route containing CBL directly followed by SRC is not, because it would require two bridges in one segment. Excluding routes that could not reappear under the rules ensures that the reproduction rate reflects competition between candidates rather than the constraints of the search. Of the admissible routes, 27 of 61 (44%) and 33 of 69 (48%) were reproduced exactly under the functional and physical STRING priors, and 100 of 117 (85%) under the regulatory STRING prior (Supplementary Fig. S7 and Supplementary Table S9).

The admissible routes that were not reproduced were short first-interval segments, such as EGFR→CBL→GRB2 and EGFR→SRC→STAT3. Once unmeasured, the deleted proteins are also no longer baseline-available, so in the first interval they compete under the degree cap *H* rather than being tested as one-step bridges independently of it (Methods). The lost routes were taken over largely by functional relatives: routes through the CBL paralog CBLB^29^ rose from 3 to 11 under the functional STRING prior and from 4 to 18 under the physical STRING prior, as did routes through GRB2 (16 to 21 and 30 to 43) and GAB1 (1 to 5 and 2 to 10), whereas these counts changed little under the regulatory STRING prior. Apart from CBL itself, the only regulatory endpoint lost was CDCP1, which had been reached by a single route, EGFR→SRC→CDCP1, that was admissible but not reproduced.

Together, these results show that, under the functional, physical, and regulatory STRING priors alike, TS-BFS-BS recovers unobserved signaling proteins largely at the positions they held when observed and retains the endpoints that depended on them. It cannot restore the anchor role of a missing protein, and it does not reproduce every route that the protein had supported.

### The order of the time intervals determines the reconstruction

The defining feature of TS-BFS-BS is that each interval starts from the proteins reached in the previous one. If the method merely connected responsive proteins through the STRING prior, the order in which the intervals are supplied would matter little. To test this, we kept the responses, the STRING prior, and all settings unchanged and supplied the three intervals of the EGF-stimulated MDA-MB-468 time course, A (0–5 min), B (5–10 min), and C (10–30 min), in two incorrect orders. In order B–C–A, the earliest responses come last; in order C–A–B, they are moved to the middle. The permuted inputs have no biological meaning; they serve to measure how much of the result is fixed by chronology.

Reordering replaced most of the reconstruction (Fig. 5a and Supplementary Table S10). Across the three STRING priors, the Jaccard indices with the correct order were only 0.23–0.33 for endpoints and 0.07–0.18 for exact routes. Proteins and edges overlapped more, but their Jaccard indices were still only 0.36–0.62 and 0.24–0.55, respectively.

**Figure 5.**
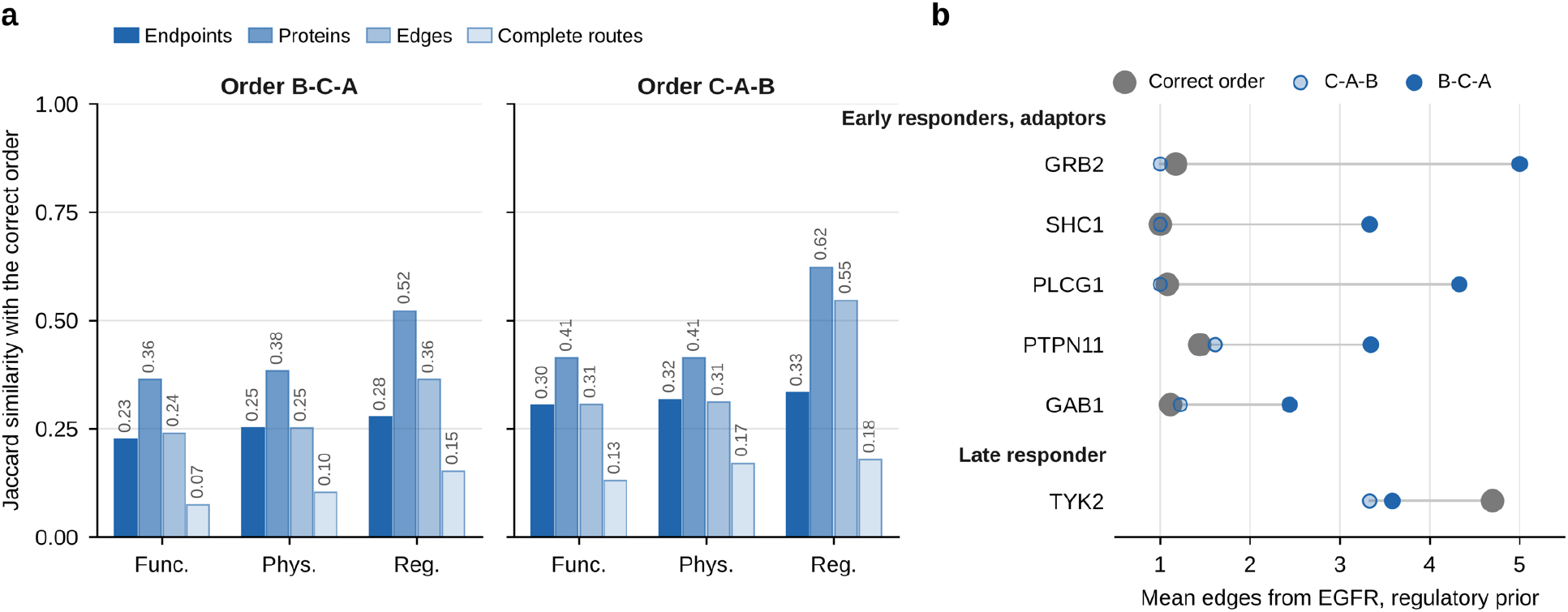
The temporal order of the intervals determines the reconstruction. The three intervals of the EGF-stimulated MDA-MB-468 time course, A (0–5 min), B (5–10 min) and C (10–30 min), were supplied in the correct order (A–B–C) or in two permuted orders (B–C–A and C–A–B), with all other inputs and settings unchanged. **(a)** Jaccard similarity of endpoints, proteins, directed edges, and complete routes between each permuted run and the correct-order run under the three STRING priors. **(b)** Mean distance from EGFR, in edges, of the receptor-proximal adaptors GRB2, SHC1, PLCG1 and PTPN11 and of the late responder TYK2 in the regulatory routes, for the correct order and the two permuted orders. See also Supplementary Fig. S8 and Supplementary Tables S10 and S11.

Because a route always ends on a protein that responds in the last interval it is given, the endpoints of each run are determined by whichever interval is placed last. In the correct order A–B–C, the last interval is C (10–30 min), so every endpoint is a protein that responds between 10 and 30 min. In order B–C–A, the last interval is A (0–5 min), and the endpoints changed accordingly: across the three STRING priors, 677 of the 680 endpoints were proteins that respond within the first 5 min. The remaining three were single routes ending on SEPTIN6 (functional and physical STRING priors) or AASDH (functional STRING prior), gene symbols that could not be matched to any measured site token in the input. Because many more proteins respond early than late (348 eligible responders in the 0–5 min interval, against 136 in the 10–30 min interval), the number of endpoints more than doubled, to 323, 285 and 72 under the functional, physical and regulatory STRING priors, against 127, 118 and 34 in the correct order. In order C–A–B, the last interval is B (5–10 min), and the endpoints were correspondingly proteins that respond between 5 and 10 min. In both permuted orders, the only endpoints shared with the correct order were proteins that respond in more than one interval and therefore remain eligible endpoints whichever interval comes last.

Few routes fit both orders. A route is admissible under a given order if it passes through one responder of each interval in that order. Only 20–35% of the B–C–A routes were admissible under the correct order, and only 30–40% of the correct-order routes under B–C–A; C–A–B gave similar figures. The routes shared with the correct order were short and ended at multi-interval responders such as CRK, PIK3R2 and TNK2, for which chronology imposes little constraint.

Placing the earliest responses last inverted the geometry of the routes (Fig. 5b and Supplementary Fig. S8). Under B–C–A, proteins that respond only early moved away from the receptor, and proteins that respond only late moved toward it. Receptor-proximal adaptors, which bind the activated receptor directly or through one adaptor^30^, became distant endpoints. Under the regulatory STRING prior, the mean distance of GRB2 from EGFR rose from 1.2 to 5.0 edges, as in EGFR→STAT3→SOCS3→TYK2→PTPN11→GRB2; SHC1 moved from 1.0 to 3.3 edges, PLCG1 from 1.1 to 4.3 and PTPN11 from 1.4 to 3.3 (Supplementary Table S11). TYK2, the most frequent regulatory endpoint in the correct order, became an intermediate in 66 routes. Under C–A–B, by contrast, the adaptors stayed next to the receptor (1.0–1.6 edges); the late responders were pulled forward instead, and STAT3 and STAT5A replaced TYK2 and STAT6 as the main regulatory endpoints.

Together, these results show that the temporal order of the measurements is a determining input rather than a formality: sequential propagation from interval to interval decides which responses become endpoints and through which proteins they are reached. Two practical consequences follow. First, the assignment of samples to intervals must be correct and documented. Second, a fraction of the routes, consisting of short routes to multi-interval responders, was produced under each of the orders tested; this fraction is the baseline against which the temporal specificity of a reconstruction should be judged.

### SHP2 inhibition and washout

We next asked whether the route ensembles follow a pharmacological perturbation. TS-BFS-BS was applied with the reference settings to two further arms of the same experiment^15^. In the SHP2i arm, cells were pretreated with the allosteric SHP2 inhibitor SHP099 (10 µM, 2 h)^31^ before EGF stimulation. In the washout arm, cells were stimulated with EGF for 10 min under SHP099, washed, and stimulated again. Each arm was run under the three STRING priors; together with the untreated runs, this gave nine ensembles (three conditions × three STRING priors), each filled to the temporal beam of 300 routes.

Two features of the design limit the comparison. First, the washout arm is a re-stimulation: its first interval spans drug exposure, the washes, and 5 min of renewed EGF, so treating EGFR as a fresh source in that interval is an approximation. Second, because the response cut-off is applied after MAD scaling within each arm (Methods), the effective threshold differs between arms (1.48-, 1.34-, and 1.38-fold for the untreated, SHP2i, and washout arms), so differences in endpoint numbers mix biology with threshold.

Condition had a larger effect than the STRING prior. Within a condition, the functional and physical STRING priors shared most of their endpoints (Jaccard 0.87–0.93), whereas between conditions under the same STRING prior the endpoint Jaccard index fell to 0.25–0.35 and the exact-route Jaccard index to 0.02–0.08 under the functional and physical STRING priors (Fig. 6a and Supplementary Table S12). Because every endpoint is a third-interval anchor in all three arms, endpoint overlap tracks the overlap of the late (10–30 min) responses (anchor-gene Jaccard 0.27–0.36), whereas the first-interval anchors are largely shared (0.62–0.70) (Supplementary Fig. S9a and Supplementary Table S13). Washout was not closer to the untreated state than SHP2i was: the endpoint Jaccard index with the untreated condition was 0.35 for washout against 0.34 for SHP2i under the functional STRING prior, and 0.35 against 0.33 under the physical STRING prior. Within 30 min, removing the drug produced a third state rather than a return to the untreated one. Under SHP2i, PTPN11 was excluded by a general eligibility rule. PTPN11 occurs in 123 untreated routes, in none of the SHP2i routes, and in 149 washout routes (of the 900 routes of each condition, three STRING priors pooled; Fig. 6b). This absence does not reflect its response to EGF, which is nearly the same in all three inputs; PTPN11 is a first-interval anchor in each. The difference lies in an additional comparison available only for the SHP2i arm. Because these cells were treated with SHP099 for 2 h before stimulation, the input also records how each site changed during pretreatment, by comparing the unstimulated SHP2i sample with the reference sample. We call this the pretreatment contrast. Each contrast site is classified with the same criterion as the interval responses: a site is resolved when its change passes the response threshold, and its function is annotated; it is opposing when it moves against the protein’s activity, that is, falls at an activating site or rises at an inhibitory one. In PTPN11, the activating sites Y546 and Y584 (Y542 and Y580 in the commonly used numbering) fell below reference during pretreatment, so both contrast sites are opposing and none is supporting. Under the eligible-pool rule (Methods), a gene whose resolved contrast sites are all opposing is treated as inactivated by the pretreatment and is excluded from every role, as anchor, bridge or source, for the whole run. PTPN11 therefore cannot appear anywhere in the SHP2i routes, whatever the STRING prior offers. Its absence is thus a consequence of the measured pretreatment contrast and of the eligibility rule, not an independent result of the search. Nineteen genes meet this criterion, including PTPN11, GAB2, MAPK1, MAPK3, PLCG2, PRKCD and SYK, as well as EGFR, which is retained as the source (Supplementary Table S14). By construction, none of the other 18 appears in any SHP2i route, whereas 227 of the 416 other anchor genes do. The rule uses measured sites only, and no gene is removed by name.

**Figure 6.**
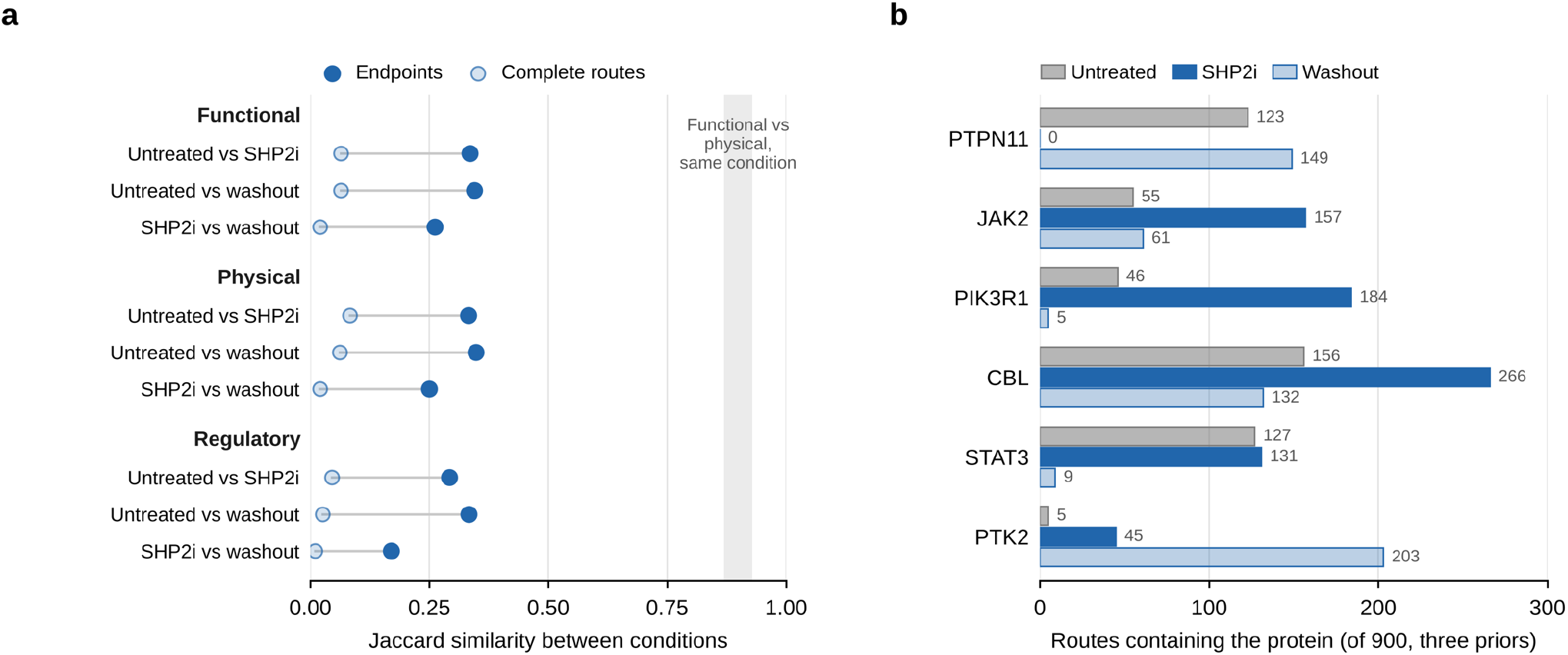
SHP2 inhibition and washout change the reached proteins more than the choice of STRING prior does. EGF-stimulated MDA-MB-468 cells were left untreated, pretreated with SHP099 (SHP2i), or subjected to inhibitor washout. **(a)** Jaccard similarity of endpoints (filled) and complete routes (open) between conditions under each STRING prior. The gray band marks the endpoint Jaccard similarity between the functional and physical STRING priors within a condition (0.87–0.93). **(b)** Number of routes containing selected proteins, pooled over the three STRING priors (900 routes per condition). PTPN11 is absent from all SHP2i routes because the eligible-pool rule excludes every gene whose resolved pretreatment-contrast sites are all opposing (Methods). Proteins that followed it in untreated routes, such as JAK2, remain reachable, mainly directly from EGFR. Under SHP2i, JAK2, PIK3R1, and CBL occur in more routes; after washout, PTPN11 returns and PTK2 rises.

What is lost under SHP2i is PTPN11 as a step, not the proteins beyond it. SRC, JAK2, STAT3 and PTK2, which remain first-interval responders in the SHP2i input, still occur in 159, 157, 131 and 45 of the 900 SHP2i routes; the search reaches them through other connections, mainly directly from EGFR, so the reconstruction is re-routed around the excluded protein. Nine of the 13 direct successors of PTPN11 in the untreated routes remain, accounting for 248 of its 272 steps, and PTPN1 becomes the first hop of 29 regulatory routes. Moreover, some excluded genes still respond to EGF under the SHP2 inhibitor (MAPK3 Y204, the GAB2 sites, and PLCG2 Y818), so their absence from the routes is not an absence of response.

Each condition had a signature that all three STRING priors reproduced (Table 4, Supplementary Fig. S9b and Supplementary Table S15). Eight endpoints were lost under SHP2i and returned after washout; 11 were specific to SHP2i, 12 to washout and 9 to the untreated condition. Across these 40 genes, a protein is an endpoint exactly when it has a non-opposing third-interval anchor and is not among the excluded genes. The categories therefore describe the 10–30 min response, not responsiveness to EGF as such. All 9 untreated-only endpoints, for example, still respond in the first washout interval, and GRB2 Y209 rises in the first interval in all three arms but falls in the third only after washout.

**Table 4.**
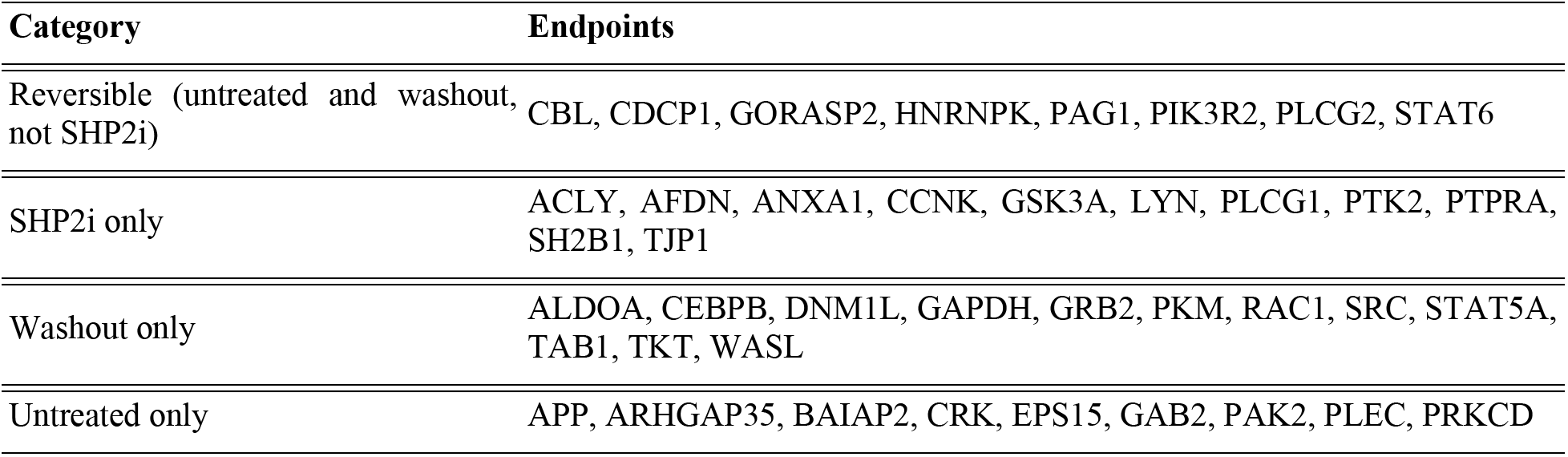
Condition-specific endpoints under SHP2 inhibition and washout compared with untreated cells. EGF-stimulated MDA-MB-468 cells were left untreated, pretreated with SHP099 (SHP2i), or subjected to inhibitor washout. Categories describe the conditions in which a protein is an endpoint; only proteins that fall in the same category under all three STRING priors are listed.

Route content shifted in opposite directions in the two arms (Supplementary Table S16). Because every ensemble holds the same number of routes, these shares describe how the beam was filled and indicate which steps the search used, not how strongly a protein is engaged. Under SHP2i, CBL became more central: its route share rose from 18.3% to 34.0% under the functional STRING prior and from 21.3% to 32.0% under the physical STRING prior. PIK3R1 occurred in 184 of 900 routes, against 46 in untreated cells, and under the regulatory STRING prior, JAK2 replaced PTPN11 as the dominant relay (44.3% of routes, against 18.3% for JAK2 in untreated cells). After washout, PTPN11 returned as the leading first hop of the regulatory run (23.0%), with outgoing edges to PTK2 (75 routes) and SRC (67). PTK2 occurred in 203 of 900 routes, against 5 in untreated cells. GRB2 became an endpoint (54 regulatory routes), whereas in the other two conditions it was a frequent intermediate but never an endpoint. PKM, ALDOA, GAPDH and TKT were washout-only endpoints under all three STRING priors, but no STRING prior provided a convincing route to them: in the functional run, 16 of the 19 routes to metabolic endpoints start at EGFR→GAPDH, a probable text-mining edge.

Several observations agree with the source study^15^. ERK1/2 and GAB2, whose key sites are below the reference after SHP099 treatment, are absent from the SHP2i routes. GRB2 Y209, identified as an SHP2 substrate in the source study, falls between 10 and 30 min only after washout, so GRB2 becomes an endpoint only in that condition. PIK3R1 gains prominence under SHP2i.

Taken together, the perturbation changes which proteins are reached far more than a change of STRING prior does. The prominent loss of PTPN11 under the inhibitor follows from measured site evidence, through a rule applied uniformly to all genes.

### Cell context: HeLa and EGFR Flp-In HEK293T with a comparison to TPS

To test whether the behavior of TS-BFS-BS depends on a single dataset, we applied the same settings to two further EGF time courses, from EGFR Flp-In HEK293T cells^6^ and HeLa cells^1^, and ran each under the three STRING priors. Because the three inputs come from different studies, they differ not only in cell line but also in experimental design and measurement. Four such differences affect the routes regardless of cell line: the type of phosphosite mainly measured, the number of time intervals, the response threshold derived from each dataset, and whether the cell-context filter could be applied (Table 5 and Supplementary Table S1). In the order MDA-MB-468, EGFR Flp-In HEK293T and HeLa: 80% of the MDA-MB-468 sites are tyrosine sites, whereas 83% and 94% of the HEK293T and HeLa sites are serine or threonine sites; the time axes comprise three, seven, and four intervals; and the response thresholds are 1.48-, 1.25-, and 1.71-fold. Finally, the expression filter was switched off for HEK293T because this line is absent from the expression resource.

**Table 5.** Three EGF time courses analyzed with TS-BFS-BS.

|  | **MDA-MB-468** | **EGFR Flp-In HEK293T** | **HeLa** |
|---|---|---|---|
| Source | Vemulapalli et al.<sup>15</sup> | Köksal et al.<sup>6</sup> | Olsen et al.<sup>1</sup> |
| Tyrosine share of phosphosite entries | 80% | 17% | 6% |
| Intervals (last interval, min) | 3 (10–30) | 7 (64–128) | 4 (10–20) |
| Response threshold | 1.48-fold | 1.25-fold | 1.71-fold |
| Expression filter | on | off | on |
| Endpoint Jaccard, functional vs physical | 0.93 | 0.93 | 0.93 |
| Route Jaccard, functional vs physical | 0.28 | 0.32 | 0.25 |
| PTPN11 in input | Y62, Y546, Y584 | S562, Y62, Y546, Y584 | not measured |
| Regulatory routes with PTPN11 | 118 (39.3%) | 124 (41.3%) | 167 (55.9%) |
| Leading regulatory module | JAK–STAT–SOCS (49.0%) | FAK–paxillin–p130Cas (74.3%) | RAS–RAF–MEK–ERK (80.3%) |

The structural behavior of the method was reproduced in all three datasets (Supplementary Table S2). All 2,697 routes (three ensembles held fewer than 300 routes) were verified to satisfy the search rules, and every endpoint is an eligible responder of the last interval. In every cell line, the functional and physical STRING priors share most of their endpoints (Jaccard ≈ 0.93) but few of their exact routes (Jaccard 0.25–0.32), and the regulatory STRING prior yields a small, concentrated ensemble whose endpoints are almost a subset of the undirected ones. Routes are longer when a time course has more intervals, but not in proportion. Under the functional STRING prior, the mean route length was 4.13 edges for the three intervals of MDA-MB-468, 4.94 edges for the four intervals of HeLa, and 6.00 edges for the seven intervals of EGFR Flp-In HEK293T; that is, 1.38, 1.24, and 0.86 edges per interval. HEK293T routes thus use fewer than one edge per interval. This is possible because a route does not have to add an edge in every interval: when a protein reached in one interval responds again in the next, the route can stay on that protein and carry it forward without a new edge. Such repeated responders are common in the long HEK293T time course, where 430 of the 593 eligible genes respond in two or more intervals.

Overlap between cell lines decayed with distance from the receptor (Fig. 7a, Supplementary Fig. S10a and Supplementary Table S17). With the STRING priors pooled, pairs of cell lines shared first hops with Jaccard indices of 0.26–0.47, proteins with 0.13–0.18, directed edges with 0.06–0.07, endpoints with 0.03–0.08, and complete routes with at most 0.011. These values are lower than those between the MDA-MB-468 input and its own interval-permuted copies (endpoints 0.23–0.33). The decay itself is largely a property of the search: the pool of high-confidence EGFR neighbors is small, and each further edge multiplies the candidates. Evidence for a shared response therefore has to come from the sites themselves and from complete routes.

**Figure 7.**
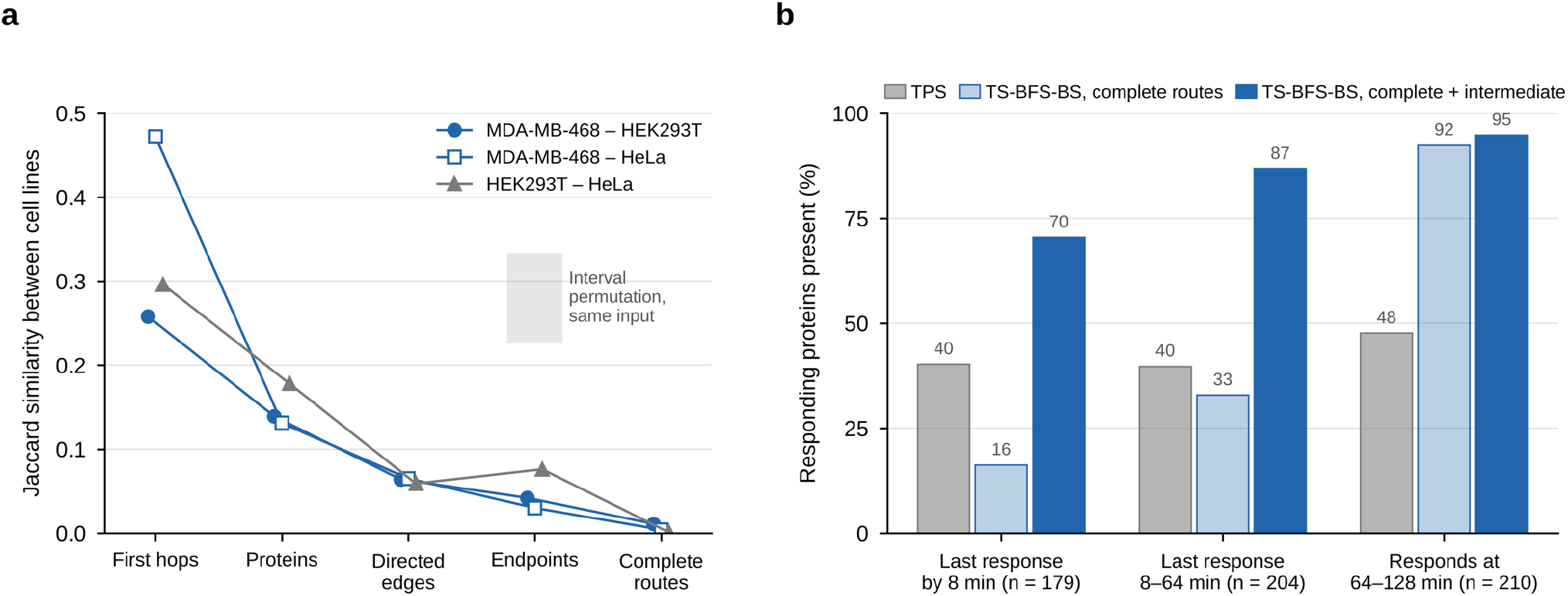
Cell context and comparison with TPS. **(a)** Jaccard similarity between pairs of cell lines (MDA-MB-468, EGFR Flp-In HEK293T and HeLa) at five levels: first hops, proteins, directed edges, endpoints, and complete routes, with the three STRING priors pooled. The gray box marks the endpoint Jaccard similarity between the MDA-MB-468 input and its interval-permuted copies (0.23–0.33). **(b)** Share of responding EGFR Flp-In HEK293T proteins present in the converted TPS network (gray), in the complete TS-BFS-BS routes (light blue) and in the complete plus intermediate routes (dark blue), grouped by the time of their last response. A protein counts as responding when one of its sites passes the 1.25-fold response threshold in an interval. Only 2,068 of the 4,113 intermediate routes are reported (Methods), so the dark blue bars are lower bounds.

At the level of sites, the early response agreed (Supplementary Fig. S10b and Supplementary Table S18). Where two experiments measured the same phosphosite, 74 of the 81 sites that respond in both first intervals changed in the same direction (55 of 60, 9 of 10 and 10 of 11 for the MDA-MB-468-HEK293T, MDA-MB-468-HeLa, and HEK293T-HeLa pairs). The concordant sites include the EGFR autophosphorylation sites, GRB2 Y209, GAB1 Y406, CRK Y136, STAT5A Y694, and the ERK1/2 activation-loop sites.

At the level of routes, MDA-MB-468 and EGFR Flp-In HEK293T shared a receptor-proximal core. The two cell lines share 16 complete routes, all running through CBL, CRK, GAB1, PTPN11, SRC, or ABL1. The EGFR→SHP2 step reappears independently in HEK293T, where the same three SHP2 tyrosines were measured and respond in the same directions (Y546 and Y584 up, Y62 down), and PTPN11 lies on 124 of the 300 regulatory routes (41.3%). In HeLa, PTPN11 was not measured, yet the regulatory STRING prior places it as a bridge between EGFR and ERK on 167 of 299 routes (55.9%) (Supplementary Table S19). In all three cell lines, PTPN11 is an intermediate and never an endpoint. This illustrates the central purpose of the method: a protein absent from the measurements is supplied where the STRING prior and the timing of its neighbors support it.

Each cell line had its own dominant modules, and each could be traced to the inputs (Supplementary Tables S16 and S20). In MDA-MB-468, these were CBL-dependent endocytosis (43–46% of undirected routes) and JAK–STAT–SOCS chains, whose endpoints, TYK2 and STAT6, are measured only in this input. In EGFR Flp-In HEK293T, they were chaperone–TP53–RNA-processing chains and, under the regulatory STRING prior, a CRK–FAK–paxillin–p130Cas module (CRK on 81.7% of routes and PTK2 on 52.7%). The regulatory endpoints of HEK293T are proximal adaptors (GRB2, CRK, CRKL and CBL) whose sites change between 64 and 128 min; they likely reflect the termination of signaling rather than its arrival at distal effectors. Without the expression filter, 5.3% of the HEK293T regulatory routes pass through T-cell receptor signaling proteins (CD247 and LAT as unmeasured bridges, and LCK, which is measured in this input; for example, EGFR→PTPN11→LCK→LAT→GRB2). In HeLa, the dominant module was an EGFR→SHP2→ERK axis (RAS–RAF–MEK–ERK in 80.3% of regulatory routes) leading to the canonical read-outs ATF2, RPS6, GSK3B, AKT1S1 and TBC1D4, whose sites are measured only in this input. Endpoint composition likewise follows the measured layer: nuclear RNA-processing and chromatin proteins make up 43% and 41% of the HEK293T and HeLa endpoints, against 12% in MDA-MB-468.

The HeLa result supports the reading that the downstream ERK and AKT–mTOR read-outs are absent from the MDA-MB-468 routes because their sites were not measured there; the same search, STRING prior and settings return these cascades once the sites are present. The routes thus differ strongly between cell lines, but every difference examined originates in what was measured, in the time axis, or in the filter settings. With these inputs, the contribution of cell context itself cannot be separated from that of the dataset.

The EGFR Flp-In HEK293T time course is the dataset for which the Temporal Pathway Synthesizer (TPS) was developed^6^, so the two methods can be compared on identical measurements. We converted the TPS network to the route format, giving a tree rooted at EGFR that reaches 309 proteins besides EGFR through 309 of the 413 published edges, one path to each protein. The published network is the union of all tree models consistent with the data, so a protein can have more than one incoming edge; a single tree keeps one incoming edge per protein, and the remaining 104 edges were left out. Besides complete routes, which span all seven intervals, TS-BFS-BS reports intermediate routes, which end in an earlier interval; this run produced 4,113, of which 2,068 are reported (at most 100 per length class; Methods).

The two methods agreed on proteins and on receptor-proximal steps (Table 6 and Supplementary Table S21). The complete TS-BFS-BS routes contain 153 TPS proteins (49.5%) and 25 TPS edges (8.1%). With the intermediate routes included, these figures rise to 244 proteins (79.0%) and 50 edges (16.2%), and nine short TPS paths are reproduced exactly. Both methods leave EGFR through CBL, GRB2, HGS, and PTK2 (Supplementary Table S22). Both also reach the nuclear serine/threonine phosphoproteins through the same unmeasured kinase, CK2α (CSNK2A1), each inferring it from its own interaction network. The disagreement on edges is mainly a disagreement between interaction networks: of 129 TPS edges leaving nine hubs of the TPS tree (EGFR, CSNK2A1, SUMO2, YWHAG, CBL, MAPK1, PRKACA, SMC3 and HDAC2; 42% of all TPS edges), 91 (71%) score below the STRING cut-off used here, so TS-BFS-BS could not have used them (Supplementary Table S23).

**Table 6.**
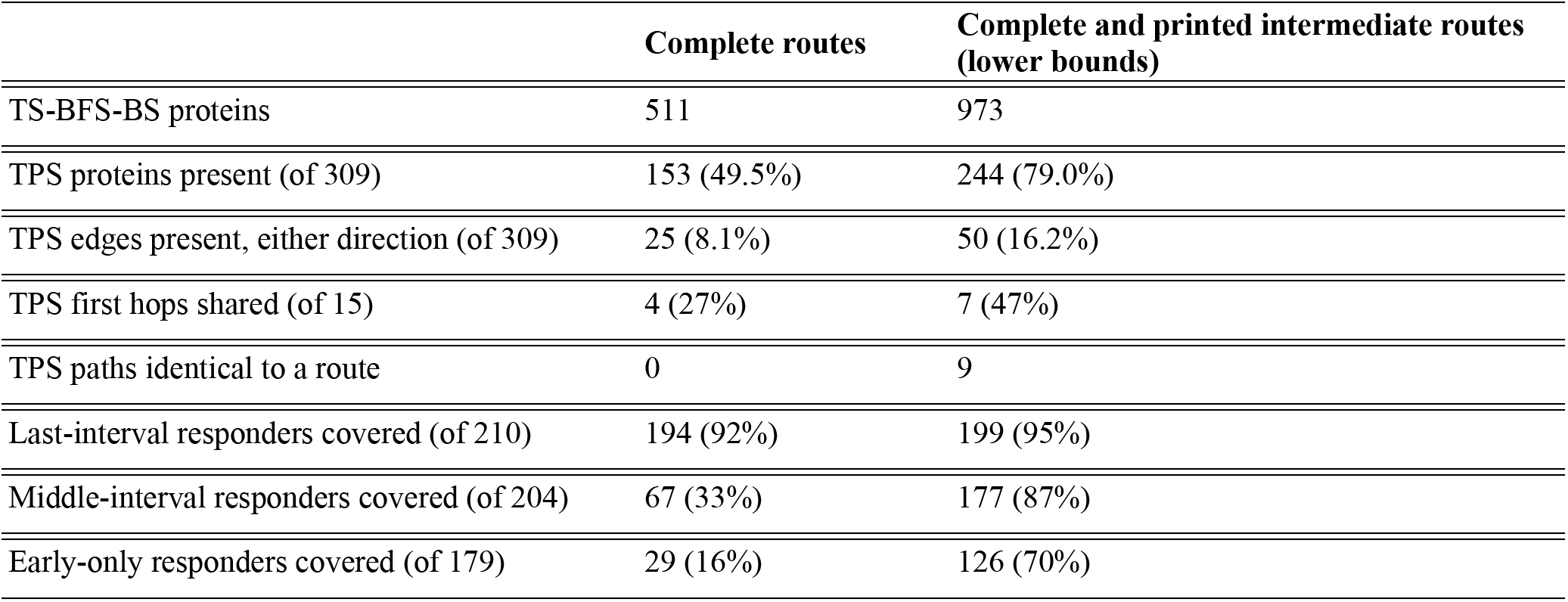
TS-BFS-BS and TPS on the same EGFR Flp-In HEK293T data. The TPS network⁶ was converted to a tree rooted at EGFR that reaches 309 proteins besides EGFR through 309 edges, one path to each protein. TS-BFS-BS results are shown for complete routes, which span all seven intervals, and with the 2,068 reported intermediate routes (of 4,113; at most 100 per length class are reported, Methods) added, so values in the second column are lower bounds. TPS edges were counted as present if the TS-BFS-BS routes contain them in either direction. Responders are grouped by the time of their last response: early (≤8 min), middle (8–64 min), and late (64–128 min).

|  | **Complete routes** | **Complete and printed intermediate routes (lower bounds)** |
|---|---|---|
| TS-BFS-BS proteins | 511 | 973 |
| TPS proteins present (of 309) | 153 (49.5%) | 244 (79.0%) |
| TPS edges present, either direction (of 309) | 25 (8.1%) | 50 (16.2%) |
| TPS first hops shared (of 15) | 4 (27%) | 7 (47%) |
| TPS paths identical to a route | 0 | 9 |
| Last-interval responders covered (of 210) | 194 (92%) | 199 (95%) |
| Middle-interval responders covered (of 204) | 67 (33%) | 177 (87%) |
| Early-only responders covered (of 179) | 29 (16%) | 126 (70%) |

The methods differ in how they treat time (Fig. 7b, Supplementary Fig. S11 and Supplementary Table S24). TPS includes 40–48% of the responding proteins in each time window, regardless of when they respond. It assigns one event to each protein, at its first significant change, and 277 of its 311 nodes are active by 8 min. When the converted TPS paths were tested against the interval rules, 136 of 309 could be laid out in temporal order, and 11 reached the last interval. The complete TS-BFS-BS routes, by contrast, reach 194 of the 210 proteins that respond between 64 and 128 min (92%), but only 16% of those that respond only within the first 8 min. With intermediate routes included, coverage is 70%, 87% and 95% for early (≤8 min), middle (8–64 min) and late (64–128 min) responders, and nine in ten TPS proteins that respond within 8 min lie on a TS-BFS-BS route. The two reconstructions are therefore complementary in time: TPS describes the spread of the initial signal, whereas TS-BFS-BS follows the response through every interval. For a long time course, the early response becomes visible only when intermediate routes are reported alongside complete ones.

The comparison does not rank the methods. TPS uses replicate statistics and accepts 203 responding proteins, whereas TS-BFS-BS applies a single fold-change threshold and accepts 602 responding genes, of which 593 are eligible as targets in at least one interval. TS-BFS-BS also relies far more on unmeasured proteins (41–47% of route proteins, against 12% for TPS), and it does not reproduce the MEK1→ERK1 order that TPS infers. Finally, its overlap with the TPS proteins is only partly specific: the complete and intermediate routes together touch 75% of all measured genes, so of the 272 measured TPS proteins, about 203 would be expected on TS-BFS-BS routes from coverage alone, against 232 observed.

## Discussion

Mass spectrometry-based phosphoproteomics is the most direct way to follow signaling as it unfolds after a stimulus, in normal cells and in disease, including the oncogenic signaling that drives cancer, because it reports thousands of site-specific changes in phosphorylation within minutes^1^. The measurement nevertheless shows only part of the system. Adaptors, scaffolds, ubiquitin ligases and other participants that act without a detectable change in phosphorylation remain hidden; enrichment chemistry, peptide detectability and stochastic sampling leave many relevant sites unobserved in any one experiment^5^; and the data alone give neither the order in which a signal passes between proteins nor the participants that link one response to the next.

TS-BFS-BS was developed to address this gap by complementing the measurements with prior knowledge. It uses STRING confidence scores, which integrate experimental, curated, co-expression, and text-mining evidence^7^. The time course gives the routes their direction, the STRING prior supplies candidate connections, including proteins that were not measured or not seen to change, and staged pruning reduces the combinatorially large space of possible routes to an ensemble small enough to inspect. The results above assess how far this approach is feasible on published EGFR data, and we discuss them in the order presented.

The parameter study showed that some outputs are much more stable than others. Endpoints, proteins, directed edges and routes of up to three edges changed little across settings, whereas longer routes were rarely reproduced. The *R × Q* grid at the degree cap used throughout (*H* = 100) made this concrete: all nine settings reached the same 127 endpoints, and 75 of the 103 reference routes of up to three edges were found in every run, but only 45 of the 197 longer routes. Runs that changed *τ*, the degree cap, the beams or the bridge depth gave the same picture: 64 of the 103 shorter routes, but only 18 of the 197 longer routes, were found in every run. This defines how the output should be read. A statement that a protein responds in a given interval and can be connected to EGFR under the STRING prior is robust, whereas a statement about one specific long route is conditional on the STRING prior and the pruning settings. Conclusions should therefore rest on the stable units, with individual routes treated as testable hypotheses rather than reconstructed mechanisms. Keeping the parameters fixed across all analyses, as done here, is what makes comparisons between conditions, controls and cell lines interpretable.

The three STRING priors encode different hypotheses rather than better or worse versions of one network^7^. They represent association, physical contact, and directed regulation, and from the same data they produced reconstructions of very different size (127, 118, and 34 endpoints; 278, 258, and 94 proteins). A reconstruction should therefore always be reported together with the STRING prior that produced it, and each STRING prior suits a different use. The functional STRING prior gave the broadest ensemble, with 127 endpoints and 92 unmeasured bridges, and reached beyond receptor-proximal signaling into endocytic trafficking, chaperones and RNA-binding proteins. It is best suited to surveying the processes to which the measured responses may connect, to generating hypotheses and to choosing read-outs for pathway-level or phenotypic experiments. Because a functional edge includes co-participation in a process, however, it implies neither contact nor direct regulation, and the prominence of hubs partly reflects how densely a process is annotated. The physical STRING prior gave an ensemble of similar breadth and behaved like the functional STRING prior in both controls. Its edges are the most direct source of candidate interactions for co-immunoprecipitation, proximity ligation or proximity labeling, starting with short segments. Physical evidence in STRING nevertheless includes membership in the same complex^7^, and physical contact establishes neither the direction nor the sign of signaling. The regulatory STRING prior was the only one to capture the SHP2 axis prominently in untreated MDA-MB-468 cells (PTPN11 on 118 of its 300 routes, against 4 and 1 under the functional and physical STRING priors), but its coverage was limited to 34 endpoints. Because its edges carry direction and sign, regulatory routes are the most suitable starting point for perturbation experiments, such as inhibitors, knockdown or knockout, and kinase–substrate assays; the SHP2 inhibitor arm is a test of this kind within the present study. The regulatory reconstruction should, however, be treated as a focused view of well-curated relations rather than a comprehensive map.

The regulatory STRING prior needs three further precautions. First, route counts per endpoint should not be interpreted. With a temporal beam of 300 and few endpoints, the beam is filled beyond the per-endpoint quota, and a few endpoints accumulate many routes (TYK2, 54); the same filling occurred under the functional STRING prior when Q was set to 1, with CRK ending with 31–38 of 300 routes. Second, a smaller temporal beam is likely to be more appropriate; we suggest roughly 30–90 routes for about 30 endpoints, that is, one to three routes per endpoint. Third, intermediates that act through new gene expression or secretion should be down-weighted or flagged; they lie on 29% of the untreated regulatory routes and 45% of those under SHP2 inhibition, and are unlikely to act within 30 min^24^. Across STRING priors, the strength of a finding should be graded by how many STRING priors reproduce it. Findings obtained under all three STRING priors, such as the backbone of 6 routes, 21 edges and 34 endpoints, and the endpoint categories of the SHP2 analysis, which are independent of the STRING prior, are the most robust because they do not depend on the semantics of any one STRING prior. Findings obtained under one STRING prior only should be attributed to the semantics of that STRING prior rather than reported as general results.

The CBL–SRC deletion tests the central premise of the method directly, namely that proteins absent from the data can be recovered from the STRING prior. With every CBL and SRC site removed from the input, the endpoint sets were almost unchanged (126 of 127, 117 of 118 and 32 of 34). Both proteins re-entered the routes as bridges, and a substantial share of the admissible original routes was reproduced: 44%, 48% and 85% under the functional, physical and regulatory STRING priors. This is the behavior the method was built for: a protein that the experiment failed to report can still be placed on a route when the STRING prior and the timing of its neighbors support it. The same behavior appeared without manipulation in HeLa cells, where PTPN11 was not measured, yet the regulatory STRING prior placed it between EGFR and ERK on more than half of the routes. The limit is equally clear. Recovery depends on the STRING prior already containing the relevant edges at high confidence, so a protein whose connections are poorly annotated, or a relation first reported in the analyzed dataset, cannot be supplied in this way. Network priors favor well-studied, multifunctional proteins, and inference that relies on them inherits this bias^32^.

The order of the measurements carries information about direction. Permuting the three intervals of the same input reduced the endpoint and exact-route Jaccard indices with the correctly ordered run to 0.23–0.33 and 0.07–0.18, and it moved receptor-proximal adaptors far from the receptor; GRB2, for instance, moved from 1.2 to 5.0 edges from EGFR. Under all three STRING priors, this effect was comparable to that of a pharmacological perturbation. The time course is therefore not a decoration of the network search: it determines which protein can follow which, and it is the source of direction in the undirected STRING priors. The assignment of samples to intervals must accordingly be documented whatever STRING prior is used. The result also argues for denser sampling in the first minutes after stimulation, when most receptor-proximal events occur^1,33^.

Perturbations changed the reached proteins more than the STRING prior did. SHP2 inhibition and washout lowered endpoint overlap with untreated cells to 0.33–0.35, against 0.87–0.93 between the functional and physical STRING priors within one condition. Washout did not restore the untreated state within 30 min but produced a third configuration in which PTPN11 returned as the leading regulatory entry point, with outgoing steps to PTK2 and SRC, and GRB2 became an endpoint. The disappearance of PTPN11 under inhibition needs to be read precisely. It follows from a general eligibility rule applied to measured pretreatment contrasts, under which every gene whose resolved sites were all opposing was excluded, and none was removed by name. Three points follow. First, what was lost was the step, not the downstream proteins: SRC, JAK2, STAT3 and PTK2 remained in the routes, and only the step through SHP2 disappeared. Second, absence from the routes is not absence of response, because some excluded genes, including ERK1, still responded to EGF under the inhibitor. Third, the changes in route content, such as PTPN1 becoming an entry point^34^ and the greater use of JAK2 and PIK3R1 under inhibition, are directed hypotheses for perturbation experiments. Whether this re-routing reflects compensatory rewiring under SHP2 inhibition^35^, for example through direct EGFR–SRC or EGFR–JAK2 coupling or through PTPN1, cannot be decided from the reconstruction; it is a hypothesis that perturbing these alternative steps under SHP099 could test. The agreement with the source study on ERK1/2, GAB2, GRB2 Y209 and PIK3R1 is encouraging^15^, but it was obtained on that study’s own data and is not an independent validation.

The behavior of the method generalized across datasets, although the biology could not yet be separated from the data. In HEK293T^6^ and HeLa cells^1^, the method showed the same structural behavior as in MDA-MB-468 cells: high endpoint agreement between STRING priors, low route agreement, and compact regulatory ensembles. Where two experiments measured the same site, the early response agreed (74 of 81 sites). Beyond the receptor, the reconstructions diverged strongly, but every difference examined could be traced to the measured layer, the time axis, or the filter settings. The downstream ERK and AKT–mTOR read-outs, for example, appeared only in HeLa, the one input in which their sites were measured. A cell-specific contribution is plausible, but demonstrating it requires matched data, or reruns that hold the measured site set, the threshold, and the expression filter fixed.

TS-BFS-BS and TPS proved complementary in time^6^. On the HEK293T data, the two methods shared the receptor-proximal steps through CBL, GRB2, HGS, and PTK2, and each, from its own interaction network, inferred the same unmeasured kinase, CK2α, as the link to nuclear phosphoproteins. They differ in how they model time. TPS assigns each protein one event at its first significant change and therefore describes the spread of the initial signal, whereas TS-BFS-BS treats each interval as a separate response; its complete routes cover 92% of the late responders, and with intermediate routes included, at least 70% of the early ones. For a long time course, the early response is therefore visible only when intermediate routes are reported alongside complete ones. The comparison does not rank the methods. TPS rests on replicate statistics, uses far fewer unmeasured proteins, and places MEK upstream of ERK in the canonical order, and most of the disagreement on edges reflects the interaction networks rather than the algorithms.

Several aspects of the current implementation limit what the results can show. Response calls rest on a robust fold-change threshold without replicate statistics, so part of what the method routes through may be measurement noise. Because the threshold is derived within each arm, the condition-specific endpoint categories of the SHP2 analysis also depend partly on the cut-off and should be confirmed at neighboring cut-offs. A bridge depth of one cannot span a multi-kinase cascade within one interval, which contributes to the sparse representation of RAS–RAF–MEK–ERK in tyrosine-dominated data. The expression filter is permissive, so tissue-restricted bridges pass it. The STRING combined scores used here include text-mining evidence, so some edges, such as EGFR→GAPDH, may reflect co-mention rather than a demonstrated interaction. Functional signs are available for only a minority of responding sites, so the polarity rules affect few genes, and signs taken from OmniPath describe the enzyme–substrate interaction rather than the site itself. A label-permutation control and a degree-preserving rewiring of the STRING prior^36^ are still needed to state how much of any overlap is specific; the temporal-order permutation tests the role of chronology but not that of the anchor identities themselves. Apart from TPS, the method was not compared quantitatively with other network-based approaches, such as prize-collecting Steiner forests^37^ or causal reasoning^11,12^, because such a comparison requires a shared benchmark with known pathways. Finally, no gold-standard pathway was available, so none of the results measure correctness. Like related prior-knowledge-guided approaches^6,12,37^, TS-BFS-BS is therefore best positioned as a generator of ranked, time-resolved hypotheses, a role that community assessments of network inference support^8^.

These limitations follow in part from a deliberate choice. By keeping the algorithm simple and free of manual intervention, we made every route traceable to a measured response and a STRING prior edge, and every difference between reconstructions attributable to the inputs. The same simplicity leaves room for noise to enter through the single threshold and for tissue-restricted bridges to pass the permissive filter. Because each component is a separate, rule-based step, it can be replaced or extended without changing the rest: replicate-based response calls, filtered or alternative STRING priors, and additional data types such as total-protein or interaction measurements can be added as the method is developed into more detailed models.

## Code and data availability

The TS-BFS-BS code, input tables, and STRING network snapshots will be made publicly available upon journal publication.

## Authors’ contributions

H.L. designed the TS-BFS-BS workflow and performed the bioinformatics analysis with the help of G.L. G.L. conceived the study and conceptualized the project, and H.L. and G.L. wrote and reviewed the manuscript.

## Ethics declarations

### Ethics approval and consent to participate

Not applicable.

### Consent for publication

Not applicable.

## Methods

The implementation follows three design principles. First, every decision is made by a general rule applied to measured evidence: responses are called by a single data-derived threshold, and genes are excluded only on the basis of their own measured sites, never by name or by the identity of a drug. Second, no curated adjustment is applied to the STRING prior; in particular, no hub blacklist was used. Third, all search settings were fixed in advance and held constant across conditions, controls, and cell lines (Table 1), so that differences between reconstructions reflect the inputs rather than tuning.

## Datasets

Three published EGF time courses were analyzed.

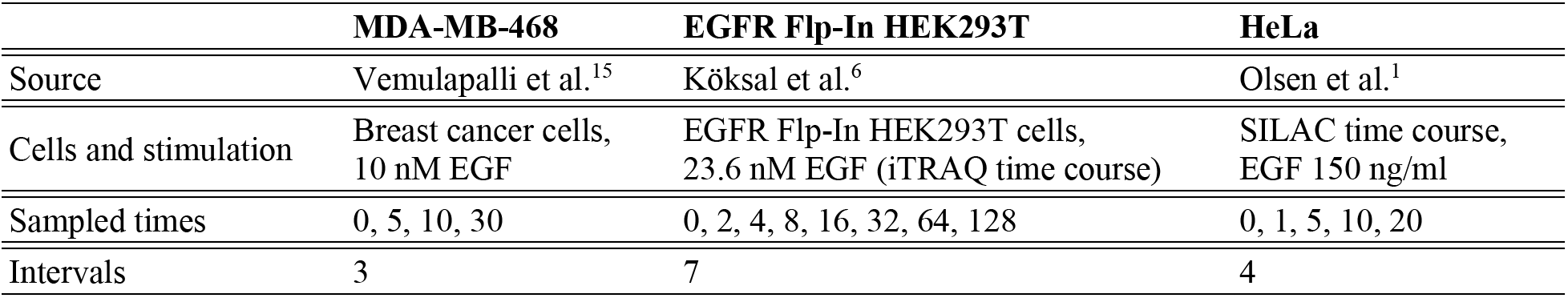

|  | **MDA-MB-468** | **EGFR Flp-In HEK293T** | **HeLa** |
|---|---|---|---|
| Source | Vemulapalli et al.<sup>15</sup> | Köksal et al.<sup>6</sup> | Olsen et al.<sup>1</sup> |
| Cells and stimulation | Breast cancer cells,<br>10 nM EGF | EGFR Flp-In HEK293T cells,<br>23.6 nM EGF (iTRAQ time course) | SILAC time course,<br>EGF 150 ng/ml |
| Sampled times | 0, 5, 10, 30 | 0, 2, 4, 8, 16, 32, 64, 128 | 0, 1, 5, 10, 20 |
| Intervals | 3 | 7 | 4 |

For MDA-MB-468, three arms were used: untreated, SHP099-pretreated (10 µM, 2 h; SHP2i), and SHP099 washout followed by re-stimulation.

Each input lists, per gene, its phosphosite tokens with normalized intensities at the sampled times. Gene symbols were harmonized to current HGNC symbols. In the SHP2i arm, the pretreatment-versus-reference contrast was stored separately and was not used as a temporal layer.

## STRING prior networks

The STRING prior is a weighted graph *G* = (*V, E, w, ξ*), where *V* is the set of proteins. *w* is the STRING v12.5 combined score, and *ξ* is the STRING prior semantics (functional, physical or regulatory). Edges below *τ* = 700 were removed (1). Neighborhoods were retrieved from the version-pinned STRING interface with the threshold applied at query time and up to 10,000 partners per protein, so that the degree cap below acts on complete high-confidence neighborhoods. Functional and physical edges are undirected; the route arrows then give only the order of traversal. Regulatory edges were traversed only from source to target, with their signs kept as annotations. Each run stored a snapshot of the neighborhoods, scores, identifier mapping, and input hash it used.

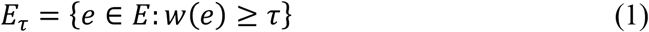

## Interval responses and anchors

The sampling times *t*_0_ < *t*_1_ < ⋯ < *t_m_* define the intervals *I_k_* = (*t_k_*_–1_, *t_k_*]. The response of token *i* in *I_k_* is computed from consecutive measurements (2), so a protein is a response anchor of *I_k_* only if it changed within that interval. Responses were scaled by a robust estimate of spread computed over all tokens and intervals of the dataset (3, 4), and anchors were selected by magnitude (5).

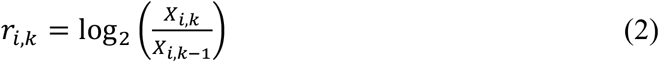

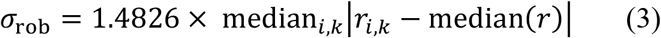

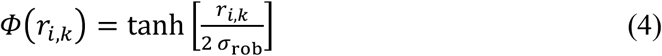

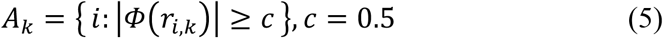

Because *σ*_rob_ is estimated per dataset, the same cut-off corresponds to a different fold change in each input (for example, 1.48-fold in untreated MDA-MB-468 and 1.25-fold in EGFR Flp-In HEK293T). The unstimulated sample *t*₀ is a reference, not an activation layer. Proteins detected at t₀ form a baseline-available set that may supply bridges in the first interval (see Sequential bounded search).

## Polarity and eligibility

Each anchor site is assigned a temporal polarity from the direction of its change and its curated functional sign *η*, which is +1 for an activating site and −1 for an inhibitory site (6). A site is classed as supporting when it increases at an activating site or decreases at an inhibitory site, as opposing in the converse cases, and as unresolved when no functional sign is available. The functional sign was assigned by exact gene–residue–position matching in UniProt canonical numbering, first from a literature-curated table of 41 regulatory sites (33 activating, 7 inhibitory and one dual-function site, EGFR Y1069) distributed with the code, and otherwise from OmniPath signed enzyme–substrate interactions (SignedPTMs)^16^, in which a site was called activating or inhibitory when the interaction between its modifying enzyme and the substrate was annotated only as stimulatory or only as inhibitory. Sites with both annotations, dual-function sites, and unannotated sites were left unresolved.

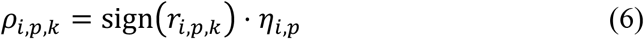

Sites were aggregated per gene. In each interval, a gene with at least one opposing and no supporting site in that interval was removed from that interval’s targets and was not admitted as a bridge in it. The gene remained available in other intervals. All other anchors, including mixed and unresolved genes, were eligible targets.

For inhibitor arms, the pretreatment-versus-reference contrast was classified with the same criterion (5), and its polarity *q* was defined as in (6). Genes whose resolved contrast sites were all opposing formed the set Q^−^ and were excluded from every role for the whole run (7). Only the stimulus source was protected from this exclusion. The rule uses measured site evidence only; no gene is excluded by name or by the identity of the drug.

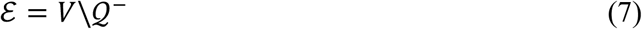

## Sequential bounded search

The search proceeds interval by interval (8). EGFR is the only source of *I*₁. The eligible anchors reached in *I_k_* become the sources of *I_k_*_+1_, so every route passes through measured anchors in chronological order. An anchor that responds again in the next interval may be carried forward without an edge (a zero-edge carry); a self-interaction is never introduced.

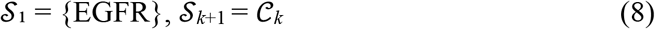

Within an interval, a bounded breadth-first search connects each source a to each eligible target b through at most *D* bridges (9). A bridge is a protein that is not an eligible target of that interval: either unmeasured, or measured without an eligible response in it. *D* counts bridges per interval and is reset at each interval, so a segment has at most *D* + 1 edges. We call the path that connects a source to a target within one interval a segment; with D = 1 it has one or two edges, and a route is the chain of segments from EGFR to an endpoint of the last interval. Target membership is tested before bridge counting, so a measured target is never consumed as a bridge, except in the first-interval baseline screen described below. Repeated proteins and immediate backtracking are disallowed. No global visited set is used, so that convergent routes through shared proteins are retained as alternatives.

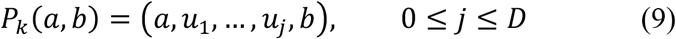

In the first interval, every baseline-available protein is additionally tested as a one-step bridge, EGFR → u → b, independently of the degree cap. A receptor neighbor that lies outside the top-ranked list can therefore still connect EGFR to an early responder. In all intervals, when the cell-context filter was enabled, a bridge was excluded only if matched RNA-seq and proteomic data for the exact cell model were both explicitly negative. Missing data were treated as unknown, and the bridge was retained, and measured anchors were never excluded on cell-context grounds. The filter was enabled for MDA-MB-468 (SIDM00628) and HeLa (SIDM00846). It was disabled for EGFR Flp-In HEK293T, which is absent from the expression resource.

## Beam pruning and hierarchical retention

Candidates are ranked by their cumulative cost (10), the sum of −ln(*w*(e)/1000) over their edges, which is equivalent to ranking by the product of the STRING combined scores. Each edge adds a non-negative cost that is smaller the higher its score, so a path ranks higher when it has fewer edges or higher-scoring edges; a longer path outranks a shorter one only when the product of its scores is larger. This ranks alternatives within the bounded search; it is not a probability of interaction. Five nested controls act on progressively larger units.

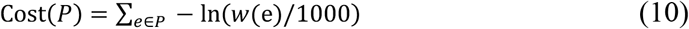

The degree cap *H* limits the expansion of each protein to its *H* highest-scoring neighbors, and the frontier beam *K_S_* limits the partial paths kept after each breadth-first layer. The segment cap *R* keeps up to *R* distinct segments per ordered source–target pair, and the history quota *Q* keeps up to *Q* upstream histories per reached target when segments are joined to the retained routes. Finally, the temporal beam *K_T_* bounds the routes propagated to the next interval. Its effective capacity is never lower than the number of reached targets (11): at least one history is kept for every reached target, then up to *Q* per target, and the remaining capacity is filled by score.

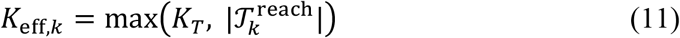

*Q* is therefore a priority quota, not a hard cap. When *Q* times the number of reached targets is below *K_T_*, as in every regulatory run here, a few well-connected targets can end more than *Q* routes. The number of routes ending at a target thus reflects how the beam was filled, not the strength of evidence.

The threshold *τ*, the degree cap *H,* and the local beam *K_S_* bound an enumeration that would otherwise grow exponentially with route length, so that the work in each interval is limited by *H*, *K_S,_* and *D*. *R, Q,* and *K_T_* allocate the retained capacity so that every reached target keeps at least one route.

The principal analysis used a STRING score threshold of *τ* = 700, a response cut-off of *c* = 0.5, at most *D* = 1 bridge per interval, a degree cap of *H* = 100, a frontier beam of *K_S_* = 300, *R* = 5 segments per source–target pair, *Q* = 3 histories per reached target and a temporal beam of *K_T_* = 300. These settings were held fixed across all analyses.

## Route assembly and output

Segments were joined only where consecutive segments shared an anchor or a zero-edge carry. Routes were collapsed by their gene chain, and phosphosite annotations were attached to genes without multiplying routes. Each node was labeled either as a measured anchor, classed as supporting, opposing or unresolved, or as a bridge, which is a protein that was unmeasured or was measured without an eligible response in that interval.

The program writes two kinds of routes. Complete routes span every interval and end on an eligible anchor of the last interval. Intermediate routes are routes as they stood at the end of an earlier interval and end on an eligible anchor of that interval; the result files print at most 100 intermediate routes per length class. The pseudocode will be made available with the code upon journal publication. Route maps (Fig. 3b and Supplementary Figs S3–S5) were drawn with a separate visualization program that annotates the retained routes but does not affect the search. Each measured anchor is drawn once per interval column and colored by the polarity of the segments that end on it in that interval: supporting, opposing, or unresolved when all such segments agree, and mixed when they carry more than one polarity class. EGFR, used only as a source, and bridges supplied by the STRING prior are drawn separately. Each edge is colored by the first of the following rules that apply. Edges graded A or B by the phosphosite check are green, and edges graded C are orange. Grade A means that the observed phosphosite position of the target matches a residue validated in the phosphosite databases (SIGNOR^38^, OmniPath^16^, iPTMnet^39^ and UniProt^40^); grade B, that the databases record phosphorylation evidence for the pair, at the residue level or as a directional interaction, but the observed site is absent or at a different position; grade C, that the pair is supported only by a non-phospho protein interaction or by UniProt phosphosites, without a confirmed kinase–substrate link; and grade D, that no database records phosphorylation evidence. Otherwise, edges to proteins classed by the cell-type check as unlikely to be expressed, or not expressed, in the cell line are red. Remaining edges take the polarity of their target anchor (green, supporting; red, opposing; purple, mixed; orange, unresolved), and edges into a bridge without such annotation are gray-blue. Edge width increases with the number of routes that use the edge.

## Statistics

For each ensemble, we counted the proteins other than EGFR, the directed edges, the route lengths, the first hops and the endpoints. We also computed the effective number of first hops and of endpoints, defined as the exponential of the Shannon entropy of their route counts, and the route share of each protein, *F*(*v*) = 100 · *N*(*v*)/*N*, where *N*(*v*) is the number of routes that contain protein *v* and *N* is the total number of routes. Route shares describe the composition of the retained ensemble; they are not estimates of abundance, activity, or flux.

Overlap between two ensembles was measured by the Jaccard index (12), computed separately for exact routes, proteins, directed edges, and endpoints. For the comparison between cell lines, first-hop usage was also compared by the cosine similarity of the frequency profiles, and protein overlap was resolved by depth, measured in edges from EGFR. All statistics are descriptive.

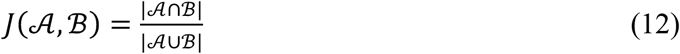

## Parameter sensitivity

Each setting was varied alone, and selected pairs were varied together, on the untreated MDA-MB-468 input under the functional STRING prior:

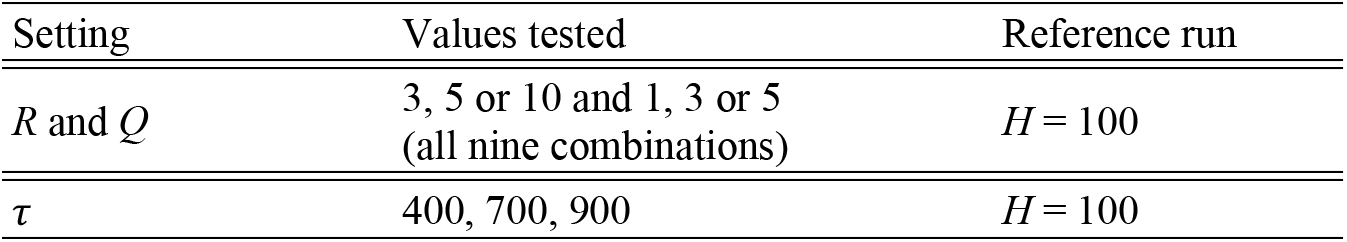
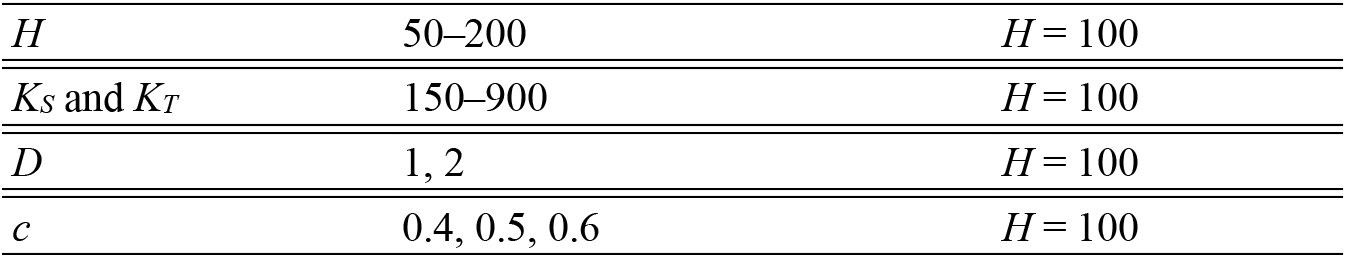

| Setting | Values tested | Reference run |
|---|---|---|
| *R* and *Q* | 3, 5 or 10 and 1, 3 or 5<br>(all nine combinations) | *H* = 100 |
| *τ* | 400, 700, 900 | *H* = 100 |
| *H* | 50–200 | *H* = 100 |
| *K<sub>S</sub>* and *K<sub>T</sub>* | 150–900 | *H* = 100 |
| *D* | 1, 2 | *H* = 100 |
| *c* | 0.4, 0.5, 0.6 | *H* = 100 |

The reference run used *H* = 100, *R* = 5 and *Q* = 3. Each run was compared with its reference by the four Jaccard indices. Route stability by length was assessed as the share of reference routes found in every run of a run set: the *R × Q* grid, or the runs that changed *τ*, *H*, the beams, or *D* (Supplementary Table S4).

## Control simulations

To test the recovery of missing proteins, we removed all CBL and SRC phosphosite entries from the untreated MDA-MB-468 input (958 to 955 tokens) and reran the three STRING priors with unchanged settings, so that the two proteins could appear only as bridges. An original route containing CBL or SRC was counted as admissible if it could still be laid out under the search rules with both proteins as non-anchors, and as reproduced if the identical gene chain occurred in the deletion run.

To test the effect of temporal order, we presented the three post-stimulation intervals of the same input to the program in the orders B–C–A and C–A–B, where A, B and C denote the correct intervals (0–5, 5–10 and 10–30 min), and left all other content unchanged. For each protein, the mean number of edges from EGFR over the routes that contain it was then compared between the correct and permuted orders.

## Comparison with TPS

The TPS network for the EGFR Flp-In HEK293T data^6^ was converted to the route format, giving one path from EGFR to each of 309 proteins and using 309 of the 413 published edges. We first counted the proteins, edges (with and without direction), and first hops shared between the TPS network and the complete and intermediate TS-BFS-BS routes. To compare coverage by response time, we divided the genes eligible by the TS-BFS-BS criterion according to the time of their last response (by 8 min, 8–64 min, or 64–128 min) and counted the share of each class present in each reconstruction. We then tested whether each TPS path could be laid out under the interval rules of TS-BFS-BS. Finally, to estimate how many TPS edges lie within the TS-BFS-BS search space at *τ* = 700, we retrieved the STRING functional scores of the 129 edges leaving nine hubs of the TPS tree (Supplementary Table S23).

## Implementation

TS-BFS-BS is implemented in Python, with modules for input normalization and identifier harmonization, response and polarity classification, construction of the eligible pool, cached retrieval of the STRING prior, bounded interval search and temporal assembly, and annotation of the resulting routes. Deterministic computations that recur across temporal histories are memoized; this shortens execution time but changes neither the search space nor the ranking. Every route in all reported ensembles was checked against its input under the search rules.

