## Supplementary Figure for "Temporal Phosphoproteomic Network Reconstruction of Condition-Dependent EGFR Signaling Routes Through Unmeasured Proteins"

### Supplementary Information Content:

**The Supplementary Information includes:**

#### Supplementary Figures and Table legends.

Legends for the Supplementary Figures and Tables, in the order of the Results. The tables are in Supplementary Data 1 (one sheet per table, with the full legend repeated above each table); every route of every ensemble is in Supplementary Data 2. Supplementary Figs S3–S6 are the route maps supplied separately.

**Contents**

| Results section | Supplementary Figures | Supplementary Tables |
|---|---|---|
| Data and runs | — | S1, S2 |
| Parameter sensitivity | S1 | S3, S4 |
| Three interaction STRING priors | S2; S3–S6 (route maps) | S5–S7 |
| CBL–SRC deletion | S7 | S8, S9 |
| Temporal-order control | S8 | S10, S11 |
| SHP2 inhibition and washout | S9 | S12–S16 |
| Cell context | S10 | S16–S20 |
| Comparison with TPS | S11 | S21–S24 |
| All routes | — | Supplementary Data 2 |

### Supplementary Figures

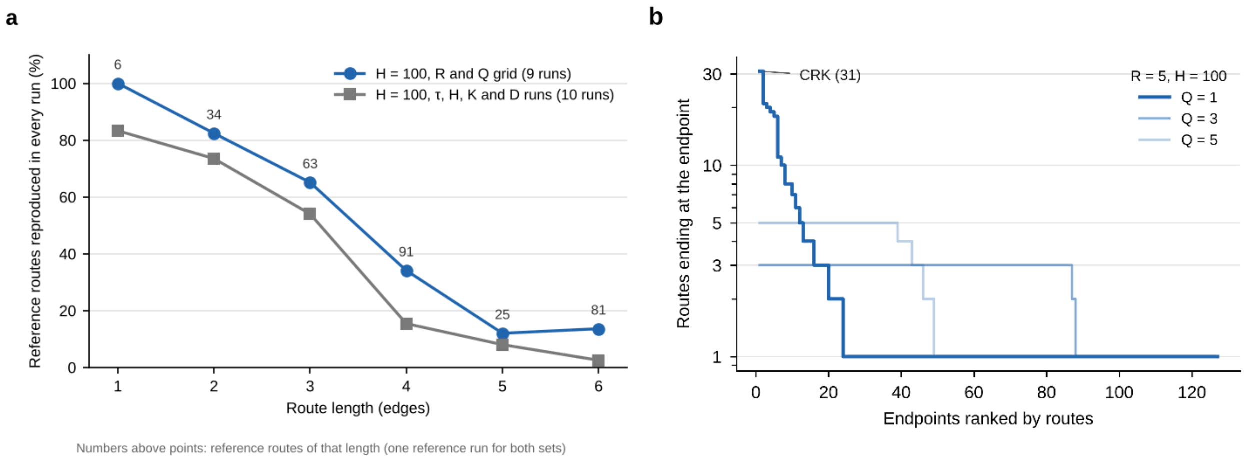


**Supplementary Figure S1 | Route stability and the history quota. (a)** Share of reference routes reproduced in every run of a run set of the parameter study, by route length. Both run sets use the EGF-stimulated MDA-MB-468 time course under the functional STRING prior and are compared with the same reference run ($H = 100$, $R = 5$, $Q = 3$). Blue, $H = 100$: the reference run and 8 runs that vary $R$ and $Q$ (9 runs). Gray, $H = 100$: the reference run and 9 runs that change τ (400, 900), the degree cap ($H = 50$, 75, 150, 200), the beams ($K = 600$) or the bridge depth ($D = 2$, and $D = 2$ with $K = 900$) without changing the input data (10 runs); the run with $K = 150$, which keeps only 150 routes, and the two cut-off runs are not included. Numbers above the points give the reference routes of that length. Routes of up to three edges were reproduced in 75 and 64 of 103 cases, and longer routes in 45 and 18 of 197, in the two run sets (Supplementary Table S4). **(b)** Routes ending at each endpoint, endpoints ranked, for $Q = 1$, 3 and 5 ($R = 5$, $H = 100$), with the same input and prior as in a. All three runs reach the same 127 endpoints. With $Q = 1$ the temporal beam is filled beyond one route per endpoint, and the surplus collects on a few hubs (CRK, 31 routes).

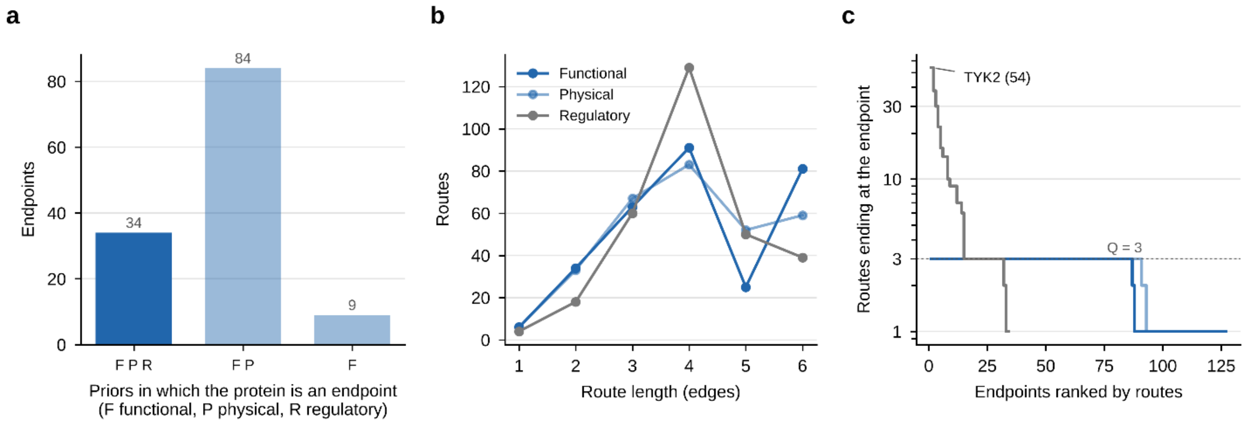


**Supplementary Figure S2 | The three STRING priors in untreated MDA-MB-468 cells. (a)** Endpoints by the STRING priors under which they are reached: 34 under all three, 84 under the functional and physical STRING priors only, and 9 under the functional STRING prior only. No endpoint is specific to the physical or regulatory STRING prior. **(b)** Number of routes of each length (edges from EGFR to the endpoint) in each 300-route ensemble. **(c)** Routes ending at each endpoint, endpoints ranked. The dashed line marks the history quota $Q = 3$. Under the undirected STRING priors, no endpoint ends more than three routes; under the regulatory STRING prior, the beam is filled beyond the quota, and TYK2 ends 54 routes. Data in Supplementary Tables S5–S7.

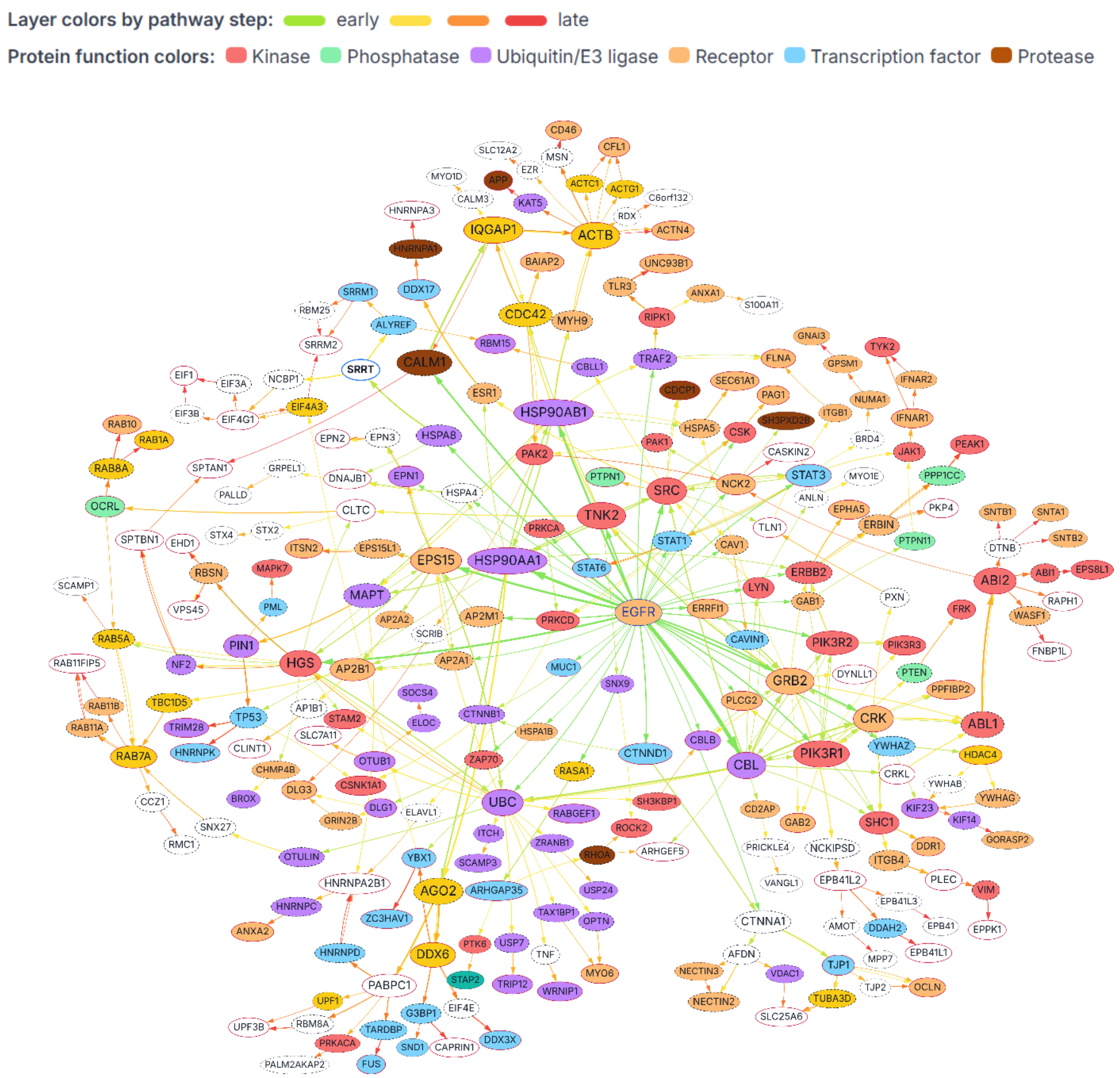


**Supplementary Figure S3 | Route network of EGF-stimulated MDA-MB-468 cells, functional STRING prior.** Union of the 300 retained routes from EGFR, with node color (protein class), solid outline (measured), and dashed outline (unmeasured bridge).

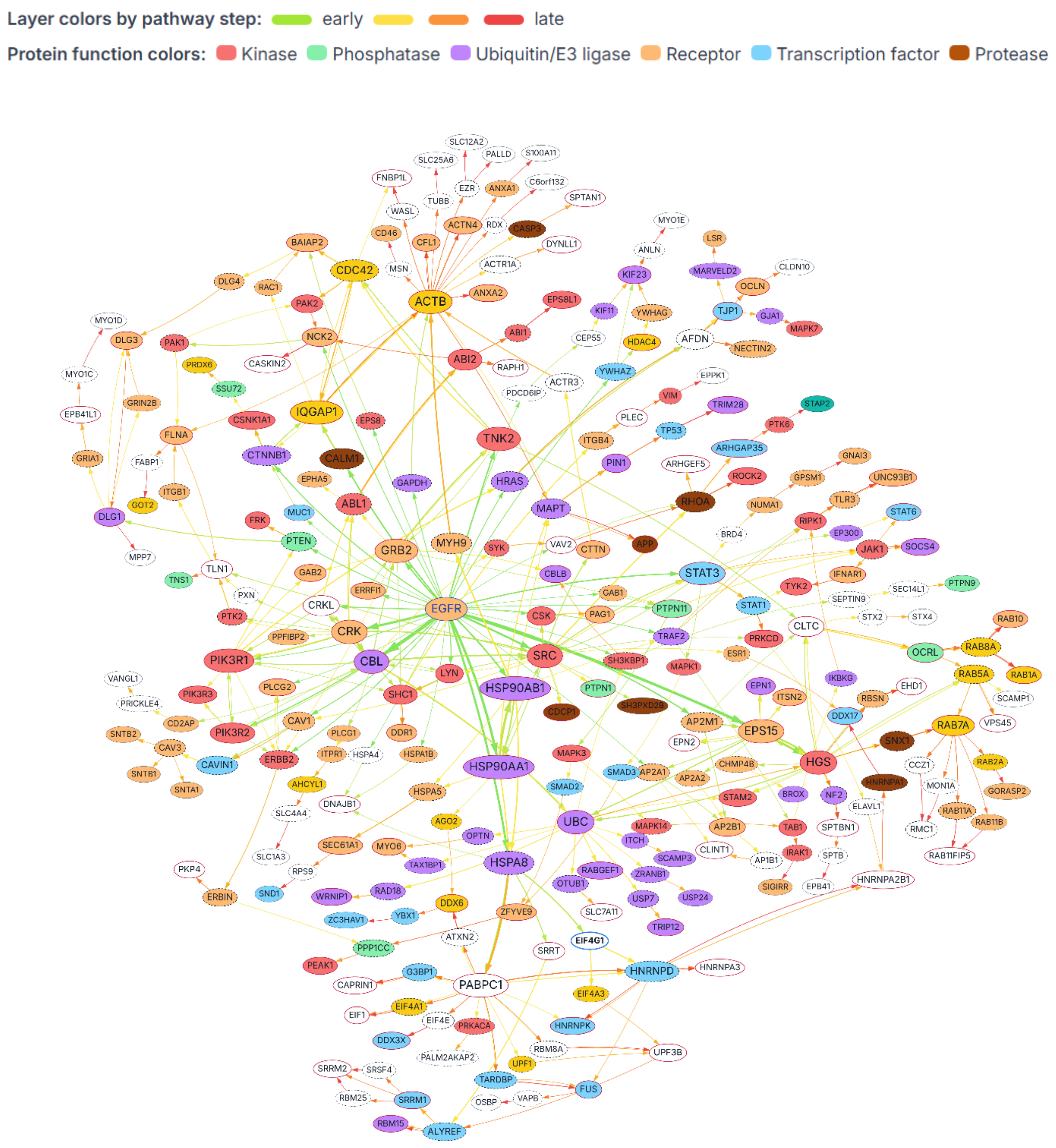


**Supplementary Figure S4 | Route network of EGF-stimulated MDA-MB-468 cells, physical STRING prior.** Union of the 300 retained routes from EGFR, with node color (protein class), solid outline (measured), and dashed outline (unmeasured bridge).

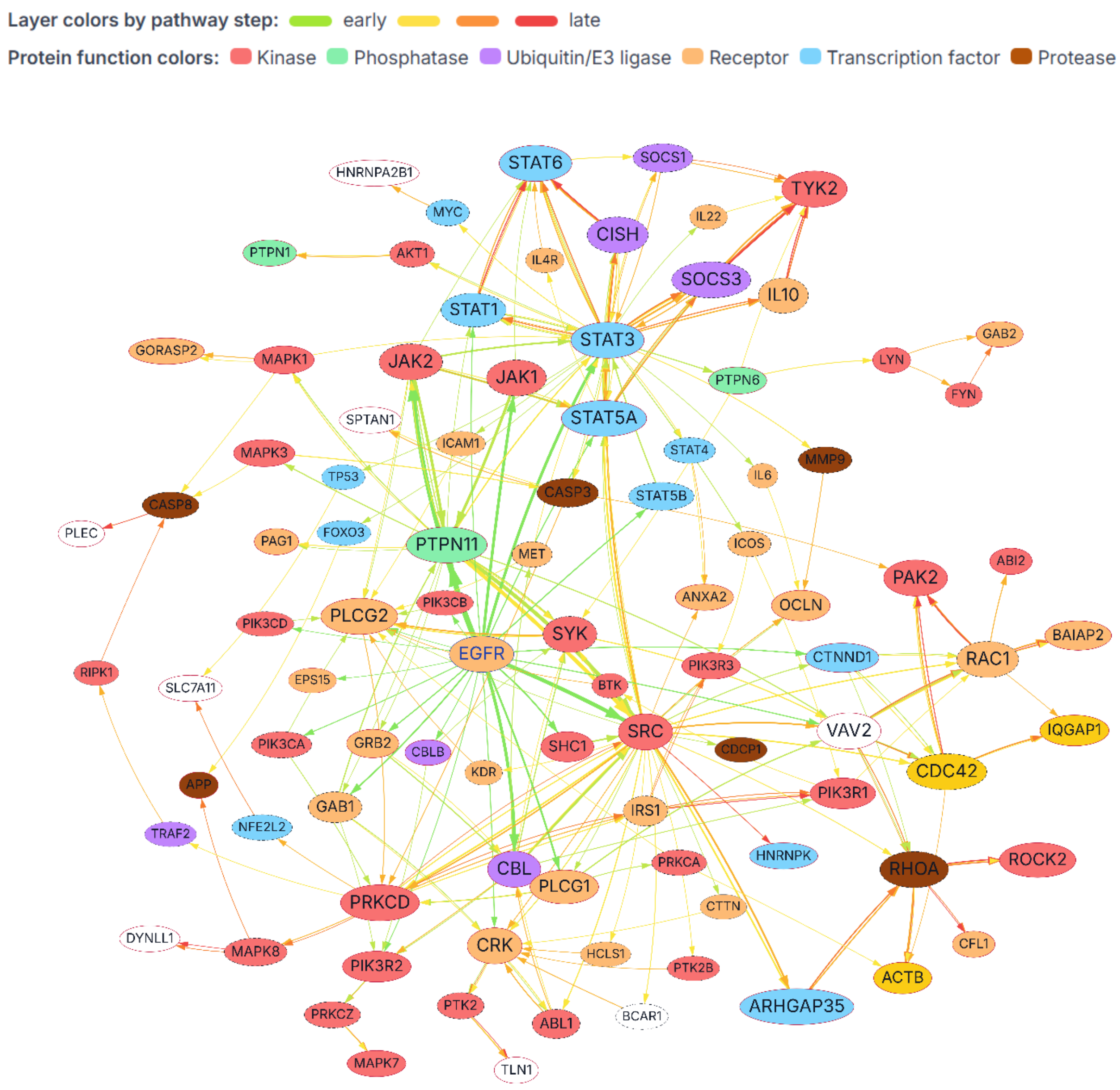


**Supplementary Figure S5 | Route network of EGF-stimulated MDA-MB-468 cells, regulatory STRING prior.** Union of the 300 retained routes from EGFR, with node color (protein class), solid outline (measured), and dashed outline (unmeasured bridge).

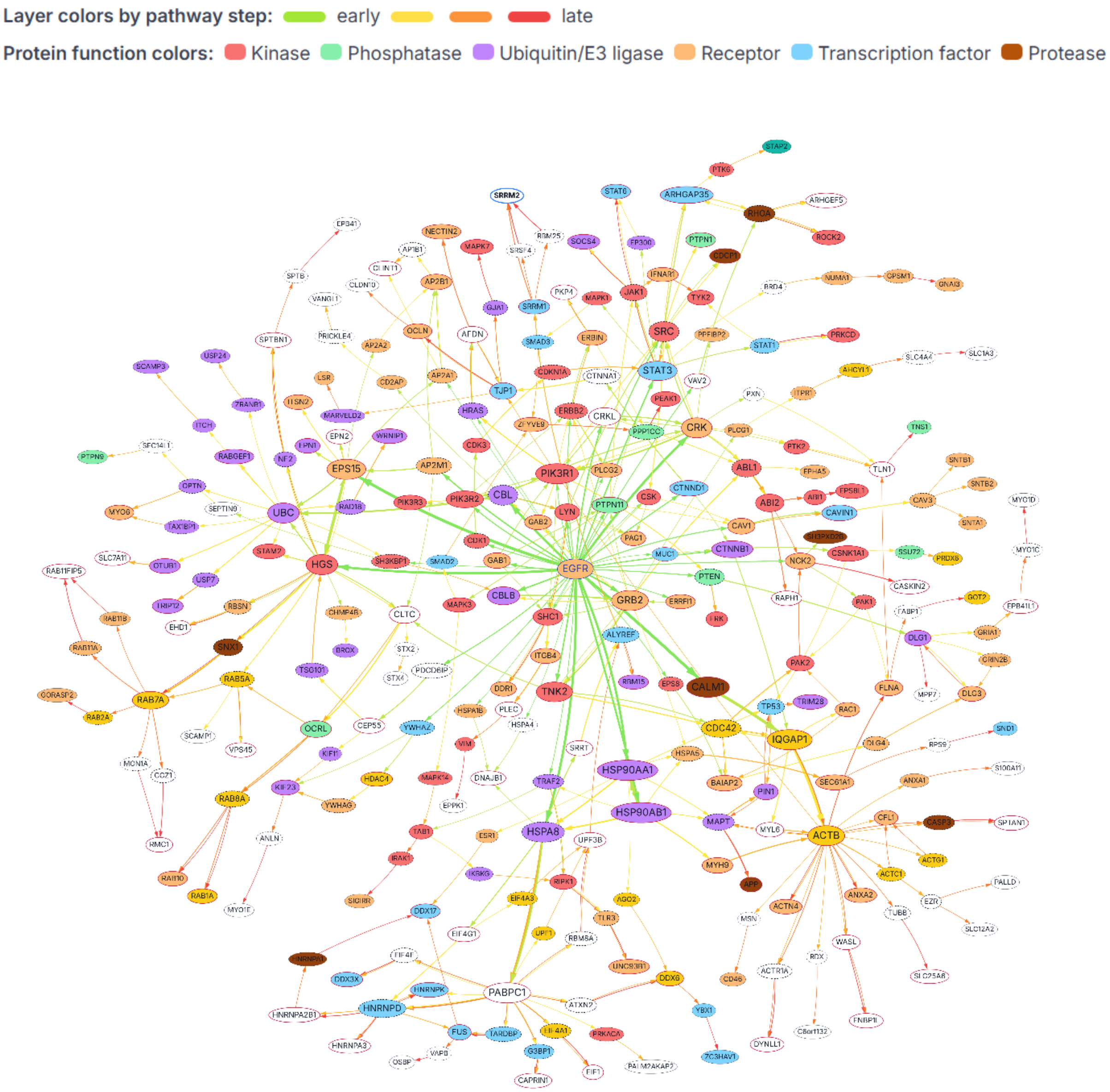


**Supplementary Figures S6 | Route network after deletion of CBL and SRC from the input, EGF-stimulated MDA-MB-468 cells, functional STRING prior.** All CBL and SRC site tokens were removed from the EGF-stimulated MDA-MB-468 input (958 to 955 tokens), and the search was rerun with unchanged settings. The network shows the union of the 300 retained routes (281 proteins other than EGFR, 405 directed edges, 126 endpoints). Solid outlines mark proteins measured in the input; dashed outlines mark unmeasured bridges supplied by the STRING prior. CBL and SRC, now unmeasured, re-entered the routes as bridges (dashed outlines): CBL in 21 routes, almost always directly after EGFR, and SRC in 18 routes, in each case between two measured anchors. Compare with the reference network in Supplementary Fig. S3.

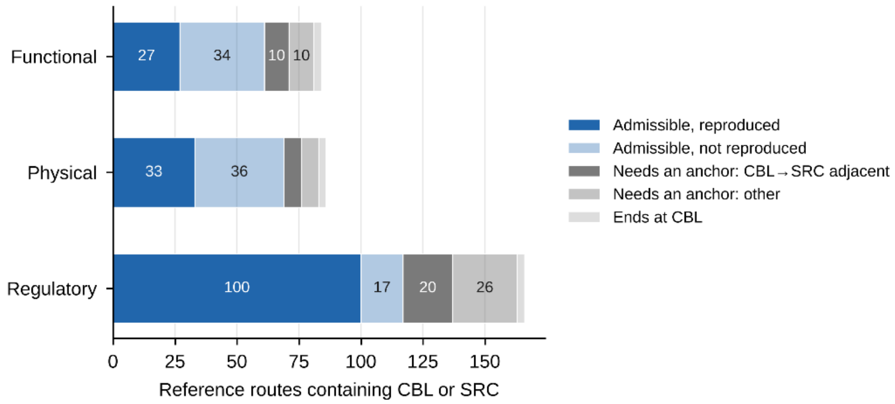


**Supplementary Figure S7 | Classification of the reference routes that contain CBL or SRC in the deletion control.** For each STRING prior, the reference routes containing CBL or SRC are divided into those that can still be laid out under the program rules once CBL and SRC can only be bridges (admissible; reproduced or not in the deletion run), those that need CBL or SRC as a measured anchor (because the two are adjacent, CBL → SRC, or for another reason), and those that end at CBL. Admissible routes were reproduced in 27 of 61 (functional), 33 of 69 (physical), and 100 of 117 (regulatory) cases. Data are in Supplementary Tables S8 and S9.

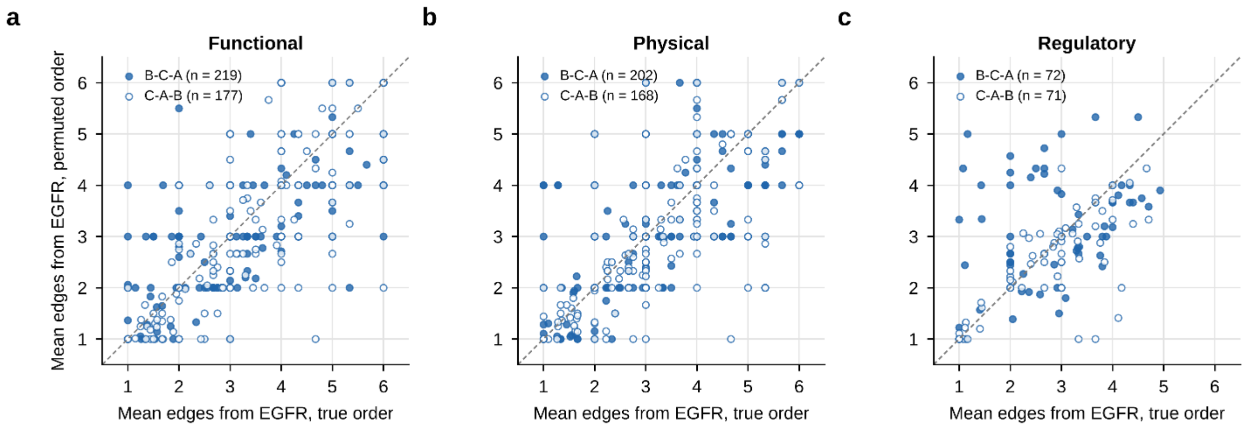


**Supplementary Figure S8 | Position of proteins in the routes under the correct and permuted interval orders.** For every protein present under both the correct order and a permuted order, the mean number of edges from EGFR over the routes containing it, under the functional **(a)**, physical **(b)**, and regulatory **(c)** STRING priors. Filled circles, order B-C-A; open circles, order C-A-B; n, number of proteins compared. The dashed line marks equal positions. Points far from the line are proteins whose place in the routes depends on the order of the measurements, such as GRB2, which moves from 1.2 to 5.0 edges from EGFR under the regulatory STRING prior. Data in Supplementary Tables S10 and S11.

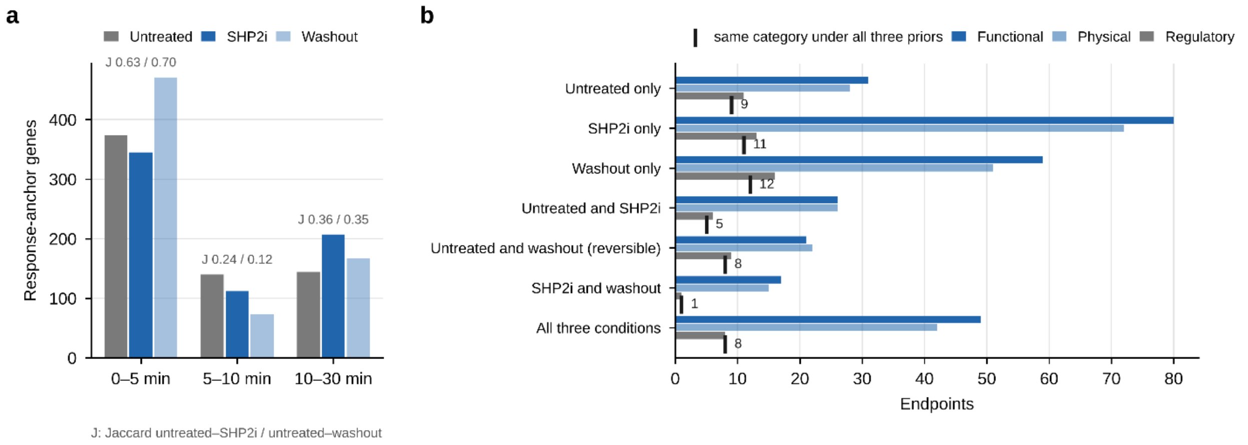


**Supplementary Figure S9 | Response anchors and endpoint categories in the SHP2 inhibition and washout conditions. (a)** Genes with a response anchor in each interval of the untreated, SHP2i, and washout inputs. J, Jaccard index between conditions (untreated–SHP2i / untreated–washout). The conditions share most first-interval anchors and diverge in the second and third intervals, which define the endpoints. **(b)** Endpoints of each STRING prior, by the conditions in which they occur. Ticks and numbers mark the endpoints that fall in the same category under all three STRING priors (STRING prior-independent endpoints; main-text Table 4). Data in Supplementary Tables S12–S15.

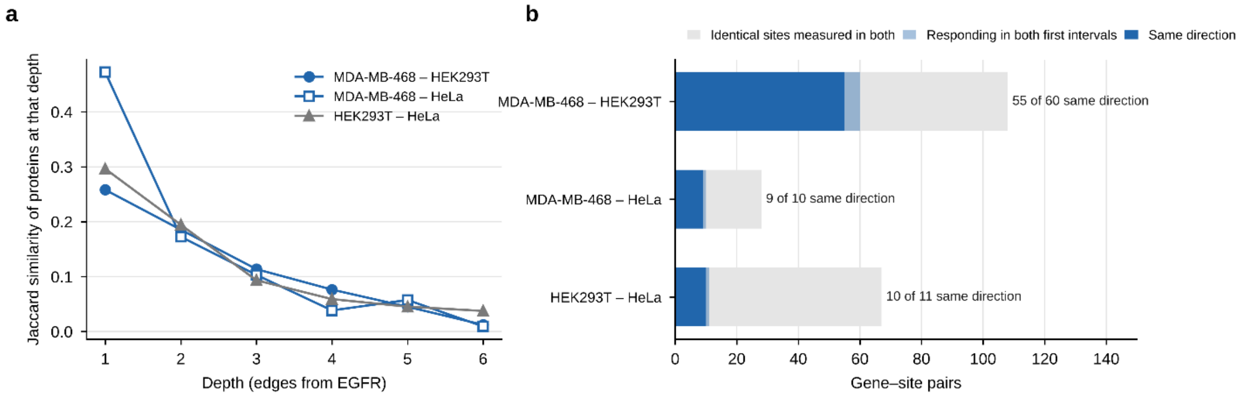


**Supplementary Figure S10 | Agreement between cell lines at the route level and at the site level. (a)** Jaccard similarity of the proteins found at the same depth (edges from EGFR) in two cell lines, three STRING priors pooled. **(b)** Identical gene–site pairs measured in two inputs (gray), those responding in the first interval of both (light blue), and those changing in the same direction (blue). 74 of the 81 sites that respond in both first intervals change in the same direction. Data in Supplementary Tables S17 and S18.

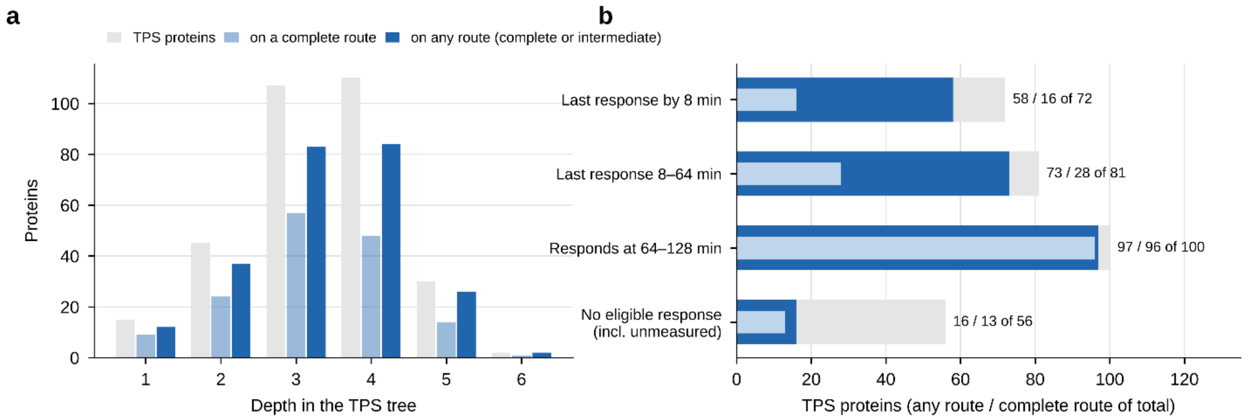


**Supplementary Figure S11 | TPS proteins in the TS-BFS-BS reconstructions of the EGFR Flp-In HEK293T time course. (a)** Proteins of the converted TPS tree by depth, and those found on a complete TS-BFS-BS route or on any printed route (complete or intermediate), with three STRING priors pooled. **(b)** The same proteins by the time of their last response (TS-BFS-BS criterion). Numbers give the proteins on any route / on a complete route, of the total. Complete routes contain most late responders; intermediate routes add most early responders. The files print about half of the intermediate routes, so the any-route counts are lower bounds. Data in Supplementary Tables S21–S24.

### Supplementary Tables (Supplementary Data 1)

**Supplementary Table S1 | Input data sets.** Source, time points, site tokens, measured genes, residue composition, robust scale, response threshold, and response-anchor and opposing-only genes per interval for the five inputs (untreated, SHP2i and washout MDA-MB-468; HEK293T; HeLa).

**Supplementary Table S2 | Route ensembles analyzed.** Size and shape of all 24 ensembles used in the paper: routes, proteins, directed edges, route length, first hops and endpoints with their effective numbers, and the largest number of routes on one endpoint.

**Supplementary Table S3 | Parameter study.** Each run compared with its reference run: endpoints, proteins, mean length, Jaccard indices for endpoints, proteins, edges and routes, reference routes retained, and endpoints lost or gained. All settings were tested against the reference run at H = 100.

**Supplementary Table S4 | Reproducibility of reference routes by route length.** Reference routes of each length reproduced in every run of the R and Q grid and of the runs that change τ, H, the beams or the bridge depth, both against the reference run at H = 100.

**Supplementary Table S5 | Agreement between the three STRING priors.** Jaccard indices with intersection/union counts for routes, proteins, edges, endpoints, and first hops between each pair of STRING priors, untreated MDA-MB-468, and the number of shared endpoints without a common route.

**Supplementary Table S6 | Protein usage in the untreated ensembles.** For each of the 350 proteins: measured status, anchor intervals, and the routes containing it, starting with it and ending at it under each STRING prior.

**Supplementary Table S7 | Backbone shared by the three STRING priors.** The 21 directed edges, 6 complete routes, and 34 endpoints found under all three STRING priors, with their route counts (main-text Fig. 3a).

**Supplementary Table S8 | CBL–SRC deletion: summary.** Endpoints, Jaccard indices with the reference run, classification of the reference routes containing CBL or SRC, and route counts of the two proteins, for each STRING prior.

**Supplementary Table S9 | CBL and SRC in the reference and deletion runs.** Route counts, identical and new routes, first-hop use, and the neighbors and positions of CBL and SRC when they enter the deletion runs as bridges.

**Supplementary Table S10 | Temporal-order control. Comparison with correct order.** Routes, endpoints, proteins, Jaccard indices with the correct order, shared routes, and endpoints lost or gained for orders B-C-A and C-A-B under each STRING prior.

**Supplementary Table S11 | Mean distance from EGFR under the correct and permuted orders.** Every protein of the nine runs with its mean position and route count under each order and STRING prior.

**Supplementary Table S12 | Overlap between untreated, SHP2i, and washout ensembles.** Jaccard indices for routes, proteins, edges, endpoints, and first hops between conditions, per STRING prior and pooled, with functional–physical agreement within each condition for reference.

**Supplementary Table S13 | Response-anchor genes per interval in the three MDA-MB-468 conditions,** with Jaccard indices between conditions.

**Supplementary Table S14 | Genes classified by the SHP2i pretreatment contrast.** The 19 opposing-only genes excluded from all roles under SHP2i (including EGFR, the protected source)

and the five mixed genes that remain eligible, with their contrast sites and route counts in the three conditions.
**Supplementary Table S15 | Endpoint categories across untreated, SHP2i, and washout.** Membership of every endpoint in the three conditions under each STRING prior, its category per STRING prior, and whether the category is the same under all three STRING priors.
**Supplementary Table S16 | Routes containing each protein in the 15 main ensembles** (three MDA-MB-468 conditions, HEK293T and HeLa, each under three STRING priors).
**Supplementary Table S17 | Overlap between the three cell lines.** Jaccard indices for first hops, proteins, edges, endpoints, and routes, and the cosine similarity of first-hop profiles, per STRING prior and pooled.
**Supplementary Table S18 | Identical phosphosites measured in two inputs.** The call of each shared gene–site pair in the first interval of both inputs and the agreement in direction.
**Supplementary Table S19 | PTPN11 (SHP2) in the three cell lines.** Measured sites, route counts, first-hop use, entry from JAK1 or JAK2, and the proteins that follow PTPN11 in the regulatory ensembles.
**Supplementary Table S20 | Module prevalence.** Share of the routes of each cell-line ensemble that contain at least one member of 17 protein modules, with the module definitions.
**Supplementary Table S21 | TPS and TS-BFS-BS on the same EGFR Flp-In HEK293T time course: summary** of shared proteins, edges, first hops and paths, without and with intermediate routes.
**Supplementary Table S22 | The 15 first-hop edges of TPS,** with their STRING score and their use in complete TS-BFS-BS routes.
**Supplementary Table S23 | STRING scores of TPS hub edges** for the 129 edges leaving nine hubs of the TPS tree.
**Supplementary Table S24 | TPS network proteins in the TS-BFS-BS reconstructions.** The 309 proteins of the converted TPS tree with depth, parent, measured status, eligible response intervals, and response class, and the complete and intermediate routes that contain them under each STRING prior.

### Supplementary Data 2

**Supplementary Data 2 | Complete routes of all ensembles.** One sheet per ensemble (24 ensembles: untreated, CBL–SRC deletion, orders B-C-A and C-A-B, SHP2i, washout, EGFR Flp-In HEK293T and HeLa, each under three STRING priors), one row per route, with route length, first hop, endpoint, and the full chain from EGFR.
